# Landscape structure drives eco-evolution in host-parasite systems

**DOI:** 10.1101/2023.10.24.563775

**Authors:** Jhelam N. Deshpande, Vasilis Dakos, Oliver Kaltz, Emanuel A. Fronhofer

## Abstract

Spatial network structure of biological systems drives ecology and evolution by distributing organisms and their genes. The ubiquitous host-parasite systems are no exception. However, past work has largely ignored relevant spatial complexity, hampering the translation of theoretical predictions to real ecosystems. Thus, we develop an eco-evolutionary metapopulation model of host-parasite dynamics where hosts and parasites disperse through realistically complex spatial networks representing major biomes: riverine aquatic and terrestrial. We generate the testable prediction that parasite virulence, or how parasites harm their hosts, is unimodal with dispersal and can reach greater values in aquatic landscapes but saturates to lower values in terrestrial systems. Moreover, we show that kin selection drives virulence evolution. Spatial networks generate characteristic patterns of parasite relatedness which drive differential virulence evolution. Finally, we show that accounting for virulence evolution allows us to predict the distribution of key epidemiological variables (e.g., parasite extinction risks) within spatial networks.

## Introduction

All biological and many artificial systems are fundamentally spatially structured. Spatial structure can be represented by networks (Urban and Keitt, 2001) where in ecology and evolution, nodes are habitat patches and links capture dispersal of organisms between these locations. Spatial network structure (topology) is key for understanding ecological patterns and processes such as the spread of perturbations (Gilarranz et al., 2017) and infections (Green et al., 2006; Davis et al., 2008; Chinazzi et al., 2020) as well as range expansions (Rayfield et al., 2023) and the distribution of biodiversity in space (Carrara et al., 2012). Importantly, spatial network structure also drives evolutionary processes: Evolution is impacted directly via gene flow within the network, but it can also be modulated indirectly via selection and drift due to the redistribution of population densities and genotypes within the spatial network (Fronhofer and Altermatt, 2017). Dispersal within the spatial network structure of landscapes is therefore a key eco-evolutionary driver (Govaert et al., 2019; Fronhofer et al., 2023).

Here, we focus on the spatial eco-evolutionary dynamics of one of the most pervasive interactions and, at the same time, maybe one of the most relevant to human and organismal health more generally: parasitism. The spatial network structure of host-parasite systems is critical for understanding epidemiological and (co)-evolutionary dynamics (Parratt et al., 2016; Penczykowski et al., 2016; White et al., 2018). For example, increased spatial clustering has been shown to reduce epidemic spread (Lilley et al., 2018), spatial network heterogeneity may lower epidemic thresholds (Colizza and Vespignani, 2008) and landscape properties impact rates of disease spread along river networks (Carraro et al., 2018). Incorporating complex landscape structures can help predict occupancy and persistence of infection (Hess, 1996): Jousimo et al. (2014) have shown in the *Plantago* (plant)-*Podosphaera* (fungal) natural host-parasite system, that increased resistance likely driven by greater gene flow in *Plantago* in highly connected habitat patches, leads to reduced parasite occupancy, when compared to less well connected patches, whereas from a purely ecological perspective higher dispersal should lead to greater parasite occupancy in more connected metapopulations (Hess, 1996).

While in epidemiology complex spatial structures have often been studied, the consequences of such spatial structures for the evolution of host-parasite systems, and how these novel evolutionary dynamics feed back on epidemiology, has not been studied as extensively. Instead, in evolutionary epidemiology, simplified spatial structures, such as lattice contact networks (Boots and Sasaki, 1999; Lion and Boots, 2010) and homogeneous (Gandon and Michalakis, 2002) or spatially implicit island models (Wild et al., 2009), have been used to understand the mechanisms driving host-parasite (co)-evolution. Previous work (Messinger and Ostling, 2009) has focused on the impact of such simplified spatial structures on virulence, a key parasite life history trait that can evolve (Frank, 1996; Cressler et al., 2016) and impact host demography (May and Anderson, 1978; Hudson et al., 1998). Virulence is defined as the extent to which parasites harm their hosts either by reducing host survival (Anderson and May, 1982) or fecundity (O’Keefe and Antonovics, 2002; Abbate et al., 2015). Parasite virulence is assumed to be a by-product of the fact that parasites need to exploit their hosts in order to transmit more (virulence-transmission trade-off hypothesis Anderson and May 1982; Alizon et al. 2009). Models of simplified spatial structures have highlighted the evolutionary mechanisms, particularly, self shading (Boots and Sasaki, 1999) and kin selection (Wild et al., 2009; Lion and Boots, 2010), that drive the evolution of parasite virulence. Generally, spatial structure can limit the availability of resources (hosts) to parasites, and also lead to genetic structuring (relatedness patterns; Lion et al. 2011) which can, in turn, lead to the evolution of reduced virulence as an altruistic strategy. Thus, while the role of spatial structure in driving the evolution of parasite virulence is widely recognised in general (Messinger and Ostling, 2009; Kerr et al., 2006), how realistically complex landscapes impact its evolution is not known.

As a proof of concept, we here consider the evolution of parasite virulence across biomes in characteristically terrestrial versus riverine aquatic landscapes. Interestingly, spatial networks, whether terrestrial or freshwater aquatic, have specific properties due to the physical processes that govern landscape genesis. Specifically, all riverine aquatic landscapes exhibit statistical properties that can be captured by optimal channel networks (OCNs) which are generated based on geomorphological processes that shape rivers globally (Rinaldo et al., 2014; Carraro et al., 2020). These networks are dendritic and and have many patches with one neighbour only, and few patches with multiple neighbours (highly heterogeneous degree distribution), which can impact, for example, metapopulation and metacommunity dynamics (Carrara et al., 2012). Similarly, terrestrial landscapes are intrinsically modular, that is, patches can be classified into modules which are highly connected within each other but sparsely connected to other modules because of biological constraints on dispersal distances. This can be captured by random-geometric graphs (RGGs; Gilarranz, 2020). Modularity has consequences, for example, for the spread of perturbations in space (Gilarranz et al., 2017).

In this context, we develop an individual-based discrete-time metapopulation model of host-parasite dynamics in which parasite virulence evolves. Spatial structure of the host-parasite system is modelled by networks representing terrestrial (random-geometric graphs; RGGs Fig. 1 A) and riverine aquatic landscapes (optimal channel networks; OCNs Fig. 1 B). Using this model we are able to track evolutionarily stable virulence in the two landscape types as a function of dispersal. We ask the following questions: 1) How does evolutionarily stable virulence differ between terrestrial and aquatic landscapes and what are the spatial network properties that drive these differences? 2) How does spatial network structure impact the evolutionary mechanisms (specifically kin selection) that drive differences in virulence evolution between terrestrial and aquatic landscapes? Finally, 3) What are the ecological consequences of evolved virulence in terrestrial vs. aquatic landscapes?

**Figure 1:**
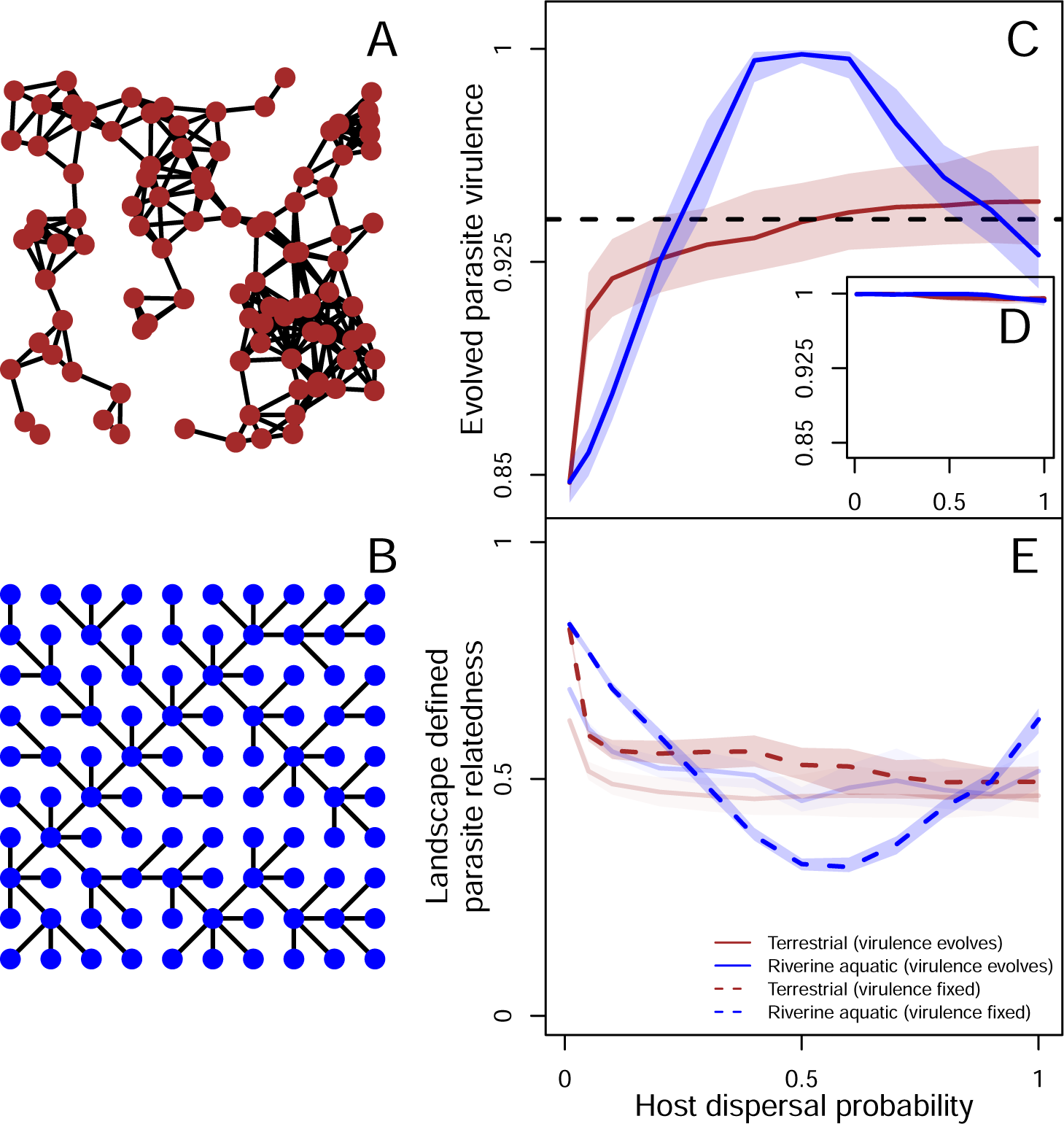
Terrestrial (A) and riverine aquatic landscapes (B) lead to differences in virulence evolution (C) which vanish when parasite kin structure is broken (D). Thus, differences in virulence evolution between the two landscape types is explained by diverging patterns of landscape defined parasite relatedness (E). A: A representative terrestrial landscape modelled as a random-geometric graph (RGG). B: A representative riverine aquatic landscape modelled as an optimal channel network (OCN). C: Evolved parasite virulence as a function of host dispersal in aquatic (blue) and terrestrial (brown) landscapes. The solid line represents the median and shaded regions the inter-quartile range of the evolved virulence trait over simulations in 1000 realisations of each landscape type, median over all infected individuals at the last simulation time step (*t* = 2500) for the simulations in which the parasite persists in the landscape. The dashed line indicates fixed virulence *v* = 0.94 which is used to represent landscape defined parasite relatedness in E. D: Evolved parasite virulence as a function of host dispersal in terrestrial (brown) and aquatic (blue) landscapes for simulations in which parasite kin structure is broken by shuffling parasites post infection, while maintaining the same infected and susceptible densities. The lines and shaded regions are the same as C, except that we include simulations where the parasite goes extinct. E: Average parasite relatedness as a function of host dispersal in terrestrial (brown) and aquatic (blue) landscapes within a patch in ecological simulations with fixed virulence (dashed lines) and when virulence can evolve (faint solid lines). We plot the average parasite relatedness over the last 100 time steps within a patch over all patches over 100 replicates. Details of landscape generation and calculation of parasite relatedness are found in the Methods and Supplementary Material. Model parameters of the focal scenario: intrinsic growth rate of the host *λ*_0_ = 4, intraspecific competition coefficient of the host *α* = 0.01, maximum parasite transmission *β_max_* = 8, shape of trade-off curve *s* = 0.5, searching efficiency of parasite *a* = 0.01.

## Methods

### Model overview

We developed an individual-based susceptible-infected (SI) metapopulation model in which parasite virulence is genetically encoded and can evolve. This model is adapted from the model developed by Chaianunporn and Hovestadt (2012). Hosts and parasites are asexual, haploid and have discrete, non-overlapping generations. Host dispersal is natal, and hosts disperse to neighbouring patches within their spatial network with a fixed probability (*d*). We assume that dispersal does not evolve, thus our model applies in cases where parasites evolve at a faster rate than hosts. Parasites rely on their hosts for dispersal. Intraspecific competition and parasite transmission are local. Virulence reduces the fecundity of the host and increasing virulence implies increasing transmission (Anderson and May, 1982; Alizon et al., 2009). We explore virulence evolution in terrestrial (represented by random-geometric graphs; RGGs Fig. 1 A) and in riverine aquatic landscapes (represented by optimal-channel networks; OCNs Fig. 1 B) of *n* = 100 patches. A summarised description of the life cycle, landscapes and analyses is given below and details are found in the Supplementary Material.

### Life cycle

#### Dispersal

After individuals are born, they disperse with probability *d*, to one of the patches connected to the natal patch with equal probability. The spatial network structure determines the connectivity of patches and is described below.

### Reproduction

After dispersal, individuals reproduce. Local population growth follows Beverton-Holt dynamics (Beverton and Holt, 1957) with intrinsic growth rate *λ*_0_ and intraspecific competition coefficient *α*:

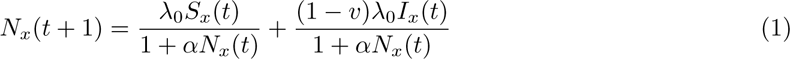

where *S_x_*(*t*), *I_x_*(*t*) and *N_x_*(*t*) are the number of susceptible, infected and total individuals in a patch *x* immediately post dispersal, *N_x_*(*t*+1) is the expected number of offspring, all of which are born susceptible. Virulence (*v*) acts on the fecundity of the host (Jaenike, 1996; O’Keefe and Antonovics, 2002; Abbate et al., 2015), so infected individuals produce on an average 1*−v* times fewer offspring relative to susceptible individuals. The realised number of offspring is Poisson distributed. After reproduction, the parental generation dies and is replaced by the offspring generation.

### Parasite transmission

Parasite spores are then released from the dead infected individuals, which then find susceptible hosts according to a modified Nicholson-Bailey model with a type-II functional response (Chaianunporn and Hovestadt, 2012; Deshpande et al., 2021), in which the transmission rate is given by *β*(*v*) and the searching efficiency by *a*:

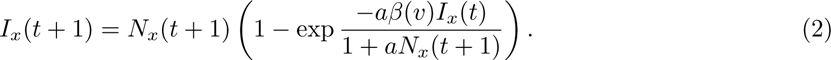

Here, *I_x_*(*t*+1) is the expected number of infected individuals in the offspring generation. The transmission rate (*β*(*v*)) of parasites is assumed to increase with virulence (Alizon et al., 2009) as:

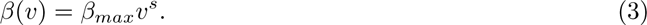

Here, *β_max_* is the maximum possible transmission rate and *s* determines the shape of the virulence-transmission relationship. Multiple parasite strains can find a host, but eventually only one can infect it. Virulence undergoes mutation with a probability *m_v_* and mutation effects on logit transformed virulence are drawn from a normal distribution with standard deviation *σ_v_*to ensure that the trait remains bounded between 0 and 1. Parasites also inherit a neutral locus which is used to track relatedness between parasite strains.

### Landscapes

We model terrestrial and aquatic landscapes using random-geometric graphs (RGGs) and optimal channel networks (OCNs), respectively. For each landscape type we generate 1000 realisations of networks of *n* = 100 patches that are encoded in adjacency matrices. RGGs are generated using the R package igraph (Csardi and Nepusz, 2006), version 3.6.3 by drawing *n* points from a *U* [0, 1] *×* [0, 1] distribution, and points that are within a radius *r* are neighbours (radius parameters of *r* = 0.15; average degree after discarding networks that are not connected; Saade et al. 2023). Increasing the radius would just lead to a metapopulation with global dispersal. OCNs are generated on an un-aggregated 10 *×* 10 grid (Carraro et al., 2020) with a fixed outlet position. The average degree of OCNs is 1.98. We do not explicitly model riverine directionality as this has been shown to only make ecological patterns stronger (Fronhofer and Altermatt, 2017).

Additionally, since terrestrial and riverine aquatic landscapes are very different in their average degree, to identify the network properties that lead to differences in virulence evolution between landscape types, we model control landscapes which are described in detail in the Supplementary Information.

### Model analysis

Our main response variable is evolved parasite virulence which we measure at the the end of each simulation (at *t* = 2500, see Fig. S1) where evolutionary dynamics have reached an equilibrium. We run simulations for all parameter combinations in Table S1, and the main text results are presented for one focal scenario that is described in all figure captions.

### Testing the role of kin selection

Previous work (O’Keefe and Antonovics, 2002; Wild et al., 2009; Lion and Boots, 2010) has argued that kin selection drives virulence evolution in spatially structured systems. This means that virulence can evolve as an altruistic strategy under conditions of limited host availability when parasite relatedness is high.

### Shuffling parasites to break kin structure

In order to test the role of kin selection in driving parasite virulence, we developed additional “shuffled” simulations in which after each transmission step, parasite genotypes are redistributed in the landscape while maintaining the same total host population size and infected densities in each patch. This breaks kin structure, without changing epidemiology (Poethke et al., 2007; Deshpande et al., 2021).

### Measuring parasite relatedness

We further test whether patterns of parasite virulence are consistent with kin selection by running additional ecological controls in which virulence is fixed (*v* = 0.8, 0.82*,…,* 1) to show how relatedness and host availability should change when virulence does not evolve. This is done in order to identify the direction of causality since, virulence is both driven by and depends on relatedness which is measured as follows:

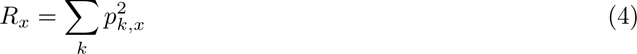

where *R_x_* is the probability of drawing two same neutral alleles in a patch *x* and *p_k,x_* is the frequency of an allele of type *k* in patch *x*. We measure relatedness within a patch because this is the scale at which parasites interact with each other.

## Results and discussion

### Landscape structure impacts the evolution of parasite virulence

#### Patterns of virulence evolution in terrestrial and riverine aquatic landscapes

We find that evolved parasite virulence differs qualitatively between terrestrial (Fig. 1 A) and riverine aquatic (Fig. 1 B) landscapes depending on host dispersal (Wild et al., 2009). For terrestrial landscapes, we observe a positive and saturating, relationship between evolved virulence and dispersal (Fig. 1 C). In contrast, aquatic landscapes produce a unimodal relationship, with peak virulence reached at intermediate dispersal values. Thus, aquatic parasites are predicted to be more virulent than terrestrial parasites for a relatively wide range of intermediate dispersal rates (Fig. 1 C; for a more comprehensive analysis, see Supplementary Material Fig. S2–S3).

Additional analyses show that these landscape type effects hold for greater maximum transmission (*β_max_*= 10) and for different shapes of the virulence-transmission relationship (saturating: s = 0.5; linear: s = 1; accelerating: *s* = 2; Fig. S2). However, for lower transmission (*β_max_* = 4) and lower host population intrinsic growth rate (*λ*_0_ = 2; Fig. S3) virulence evolution depends only on host dispersal and not on landscape structure. This is likely because in our model, high levels of virulence and transmission lead to strong oscillations in host-parasite densities (Deshpande et al., 2021). Going out of this dynamical regime (see Fig. S4) reduces the differences between the two landscapes because lower amplitude oscillations do not induce local extinctions of the host. Thus host availability is not limited and parasite kin selection becomes less important (see below).

### Which network properties explain parasite virulence evolution?

Since the processes that generate terrestrial (Gilarranz, 2020) and aquatic (Rinaldo et al., 2014) landscapes are fundamentally different, these networks differ in a range of properties, including their average degree, that is, the average number of links between patches (see Fig. 2 A). Using a series of control landscapes where we can fix the average degree while varying other properties (see Fig. S5 and S6 for details) we show that the results presented in Fig. 1 C are not due to mere differences in the average number of links. Rather, virulence evolution in OCNs is mainly driven by the presence of large numbers of headwaters, that is, patches that are only connected to one other patch (Fig. 2 A; Fig. S5). Our results for RGGs critically depend on networks being spatially embedded (as opposed to networks such as random graphs which do not have a spatial generating process) and, to a lesser degree, being modular (Fig. S6).

**Figure 2:**
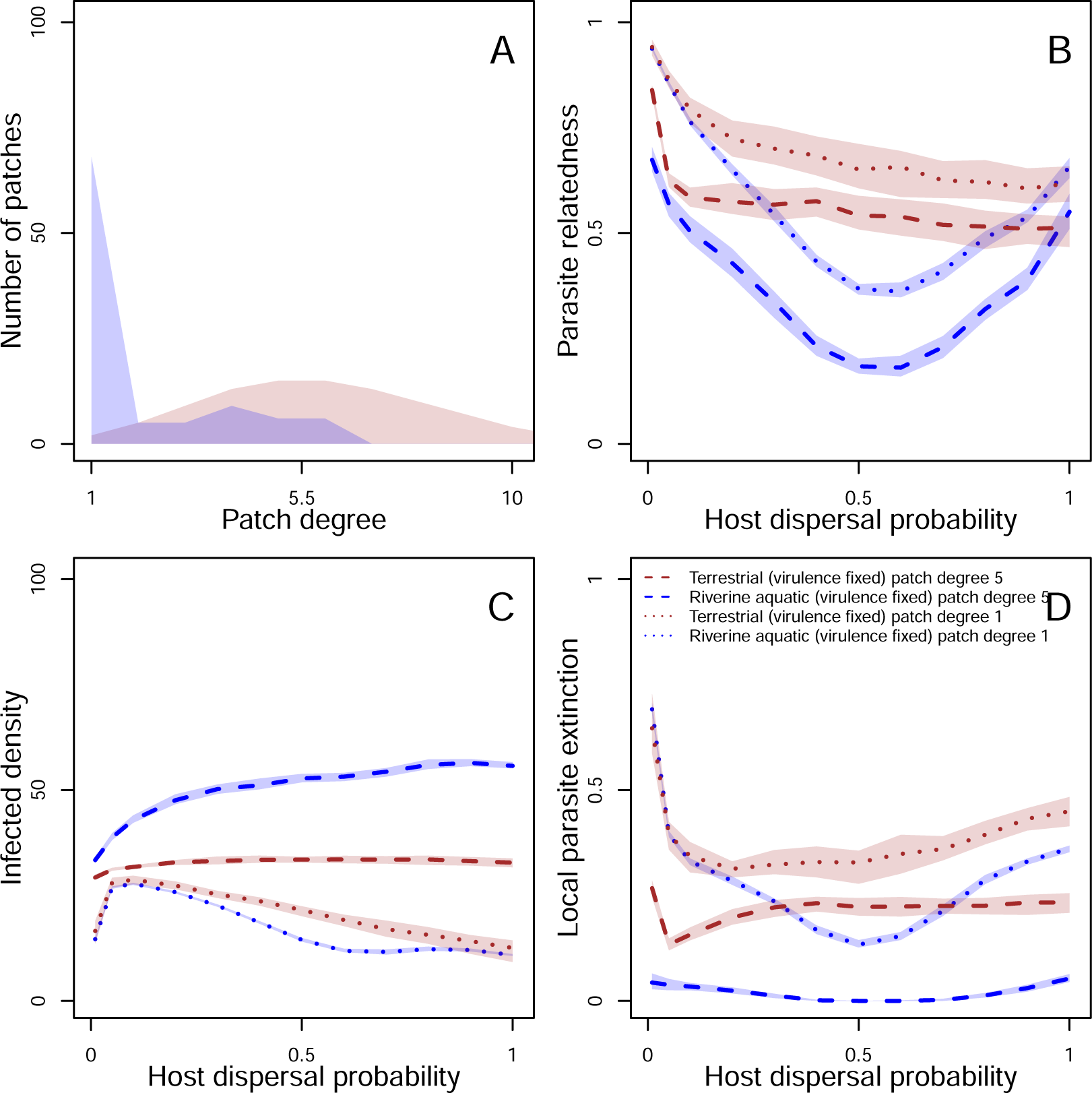
Degree distribution (A) and parasite relatedness (B), infected densities (C) and local parasite extinctions (D) as a function of dispersal for low (degree-1) and high (degree-5) connectivity patches for riverine aquatic and terrestrial landscapes. A: Degree distribution of terrestrial vs. riverine aquatic landscapes. The x-axis represents patch degree and the shaded region shows the median frequency of a patch of a given degree over 1000 landscape realisations for both landscape types. B: Parasite relatedness as function of dispersal rate for low and high degree patches. C: Average density of infected individuals for low and high connectivity patches in the landscape as a function of host dispersal over the last 100 time steps. D: Local parasite extinction calculated as the fraction of time over the last 100 time steps for which the parasite is extinct for low and high connectivity patches. For all plots, medians are taken over for 100 and 1000 realisations for ecological and evolutionary simulations respectively. Model parameters of the focal scenario: intrinsic growth rate of the host *λ*_0_ = 4, intraspecific competition coefficient of the host *α* = 0.01, maximum parasite transmission *β_max_* = 8, shape of trade-off curve *s* = 0.5, searching efficiency of parasite *a* = 0.01.

### Landscape structure drives parasite virulence evolution via kin selection

Clearly, network topology is an important driver of parasite virulence evolution. But what are the relevant evolutionary mechanisms and how are they modulated by landscape structure?

### Kin selection drives differences in virulence evolution between terrestrial and riverine aquatic landscapes

We show in Fig. 1 D that when kin structure is broken by shuffling parasites at every generation (Poethke et al., 2007; Deshpande et al., 2021), virulence evolves to its maximal value (*v* = 1) in both terrestrial and riverine aquatic landscapes. Thus, kin selection drives differences in virulence evolution between the two landscape types. More specifically, maximal virulence evolves as expected (O’Keefe and Antonovics, 2002; Abbate et al., 2015) in both landscape types when kin structure is broken because increase in virulence leads to increased transmission, without an individual cost to parasite fitness which is a feature of models of virulence acting on fecundity (O’Keefe and Antonovics, 2002). However, when kin structure is intact (Fig. 1 C), locally parasites are related to each other (that is, they share identical alleles), and under conditions of limited host availability (Fig. S7) virulence may evolve to values that are below their maximal possible value, and more importantly may depend on landscape structure (Fig. 1 C). Thus, our result is consistent previous work on evolution of virulence acting on host fecundity (O’Keefe and Antonovics, 2002) and mortality (Wild et al., 2009; Lion and Boots, 2010) that highlights the central role of kin selection in driving parasite virulence evolution in spatially structured host-parasite systems (O’Keefe and Antonovics, 2002; Wild et al., 2009; Lion and Boots, 2010). Below we show how landscape structure and host dispersal modulate parasite relatedness.

### Patterns of virulence evolution mirror within-patch relatedness in terrestrial vs. aquatic landscapes

Therefore, since differences in virulence evolution between the two landscape types are driven by kin selection, we would expect that more virulent parasites evolve when local parasite relatedness is low and vice versa, given that host availability is similarly limited between the two landscape types (Fig. S7). Indeed, Fig. 1 E shows that riverine landscapes produce a U-shaped relationship between relatedness and dispersal attaining minimum relatedness lower than terrestrial landscapes at intermediate dispersal rates, whereas terrestrial landscapes produce a saturating decrease. These patterns exactly mirror the relationship between dispersal and evolved virulence (Fig. 1 C). Note that parasite relatedness and host availability are measured in simulations in which virulence is fixed (*v* = 0.94 in the main text, a wider range of virulence values is found in Fig. S7) since virulence both impacts relatedness (Fig. S7) and is driven by it (Fig. 1 D). The mirroring of parasite relatedness and evolved virulence also holds for all control landscapes (Fig. S8–S9). Thus, ultimately, differences in how landscapes structure genotypes in space, as captured by parasite relatedness (Fig. 1 E) drives diverging patterns of virulence evolution (Fig. 1 D). Below, we explain the spatial network properties that lead to differences in parasite relatedness, hence virulence evolution between terrestrial and riverine aquatic landscapes.

### Spatial network properties determining parasite relatedness

We now seek to understand the network properties that drive differences in within-patch relatedness between terrestrial and riverine aquatic landscapes. As shown above, the shape of the degree distribution is relevant, particularly the large amount of degree-1 patches in aquatic landscapes (Fig. 2 A). For both landscapes, parasite relatedness is low in patches with high connectivity and high in patches with low connectivity (Fig. 2 B). Further, at lower dispersal rates (up to *d* = 0.1), local connectivity dominates and all landscapes have similar distributions of parasite relatedness given a patch degree. But since there are many more degree-1 patches in riverine aquatic landscapes (Fig. S10), overall relatedness averaged over all patches in the landscape is greater. As dispersal increases, due to the heterogeneity in degree distribution in riverine aquatic networks (specifically, many patches of degree-1 connected to few patches of larger degree and few patches of a larger degree connected to many patches of degree-1), there is greater dispersal out of low-connectivity patches and lower dispersal into them (Fig. S11; Fronhofer and Altermatt 2017; Altermatt and Fronhofer 2018), which leads to relatively lower infected densities in low degree patches and larger infected densities in high degree patches relative to terrestrial networks (Fig. 2 C). This implies that the parasite rarely goes extinct in high degree patches and recolonises low degree patches of aquatic landscapes, reducing the effect of genetic drift in the landscape and leading to lower relatedness relative to terrestrial landscapes (Fig. 2 D). Finally, as dispersal increases further, infected densities are further reduced in low degree patches, which leads to increasing extinction in high degree patches, and overall greater relatedness.

These analyses of neutral genetic diversity provide a mechanistic explanation of how landscape network structure shapes parasite relatedness, through the specific patterns of patch connectivity and the concomitant dynamics of patch extinction and (re)-colonisation.

### Closing the eco-evolutionary feedback loop: Ecological consequences of virulence evolution in aquatic and terrestrial systems

While the above sections worked out the mechanisms driving virulence evolution, here we ask how evolved parasite virulence feeds back on spatial epidemiological dynamics. We focus on a key epidemiological variable, local parasite extinction, i.e., the fraction of time there is no parasite in a given patch. As shown above (Fig. 2 D), for purely ecological scenarios, there is characteristic variation in extinction risk according to network type. Below, we will assess the additional impact resulting from virulence evolution, for different levels of host dispersal. As a general rule, except for very low dispersal rates (see Supplementary Material Fig. S12 and Fig. S13), virulence is independent of patch connectivity (degree; Fig. 3 A–C). Thus, differences in virulence evolution between landscape types (Fig. 1 C), and not within a landscape (Fig. 3 A–C) should explain differences in local parasite extinction patterns that will be described below.

**Figure 3:**
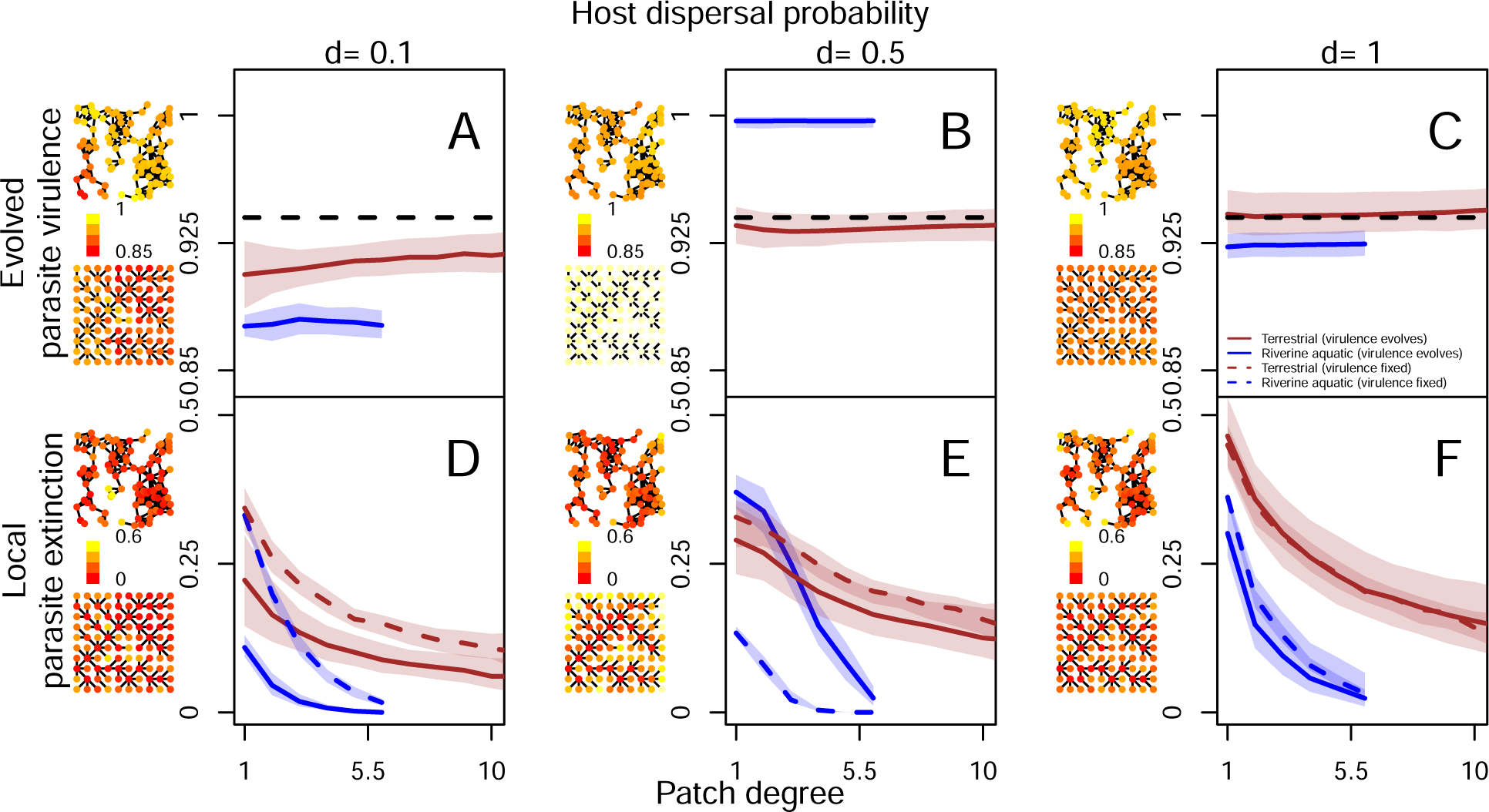
Spatial distribution of evolved parasite virulence (A–C) and local parasite extinction (D–F) as a function of patch degree for terrestrial (brown) and aquatic (blue) landscapes for different host dispersal probabilities (*d* = 0.1, 0.5, 1) in ecological (virulence *v* = 0.94, dashed line) and evolutionary (solid lines) simulations. From left to right, dispersal probability increases. A–C: Evolved parasite virulence in a patch, averaged over the last 100 time steps is plotted as a function of patch degree. On the left side of each plot, the distribution of evolved virulence averaged over the last 100 time steps for one representative terrestrial and aquatic landscape is shown with more yellow colours indicating higher virulence. D–F: Local parasite extinction calculated as the average fraction of time in which parasite is absent from a patch is plotted as a function of patch degree. On the left side of each plot, the distribution of local parasite extinction over the last 100 time steps for one representative terrestrial and aquatic landscape is shown with more yellow colours indicating higher local parasite extinction. For all plots, medians are taken over for 100 and 1000 realisations for ecological and evolutionary simulations respectively. Model parameters of the focal scenario: intrinsic growth rate of the host *λ*_0_ = 4, intraspecific competition coefficient of the host *α* = 0.01, maximum parasite transmission *β_max_* = 8, shape of trade-off curve *s* = 0.5, searching efficiency of parasite *a* = 0.01.

Without accounting for differences in parasite virulence in both landscape types, thus in simulations in which virulence is fixed (*v* = 0.94 in the main text for illustrative purposes; see Supplementary Material Fig. S12 and Fig. S13 for a wider range of virulence values), terrestrial and aquatic landscapes produce distinct relationships between patch connectivity and local parasite extinction probability (Fig. 3 D–F). Depending on the level of evolved virulence, extinction risk is shifted towards higher or lower levels, relative to this ecological reference scenario of fixed virulence (Fig. 3 D–F).

At relatively low dispersal rates (*d* = 0.1; Fig. 3 D), from a purely ecological perspective, low connectivity patches have similar high local parasite extinction for a fixed equal virulence in terrestrial and aquatic landscapes. But evolution of lower virulence in riverine aquatic landscapes leads to reduced local parasite extinction irrespective of connectivity relative to terrestrial landscapes. At intermediate dispersal (*d* = 0.5; Fig. 3 E), without accounting for differences in virulence evolution, one would predict lower parasite extinction as a function of connectivity throughout the landscape in riverine aquatic relative to terrestrial landscapes due to the heterogeneity in degree distribution (Fig. 2 A) maintaining high infected densities (Fig. 2 C) such that all patches in the landscape are always recolonised. However, the evolution of greater virulence in riverine aquatic landscapes means that local parasite extinction is higher in low connectivity patches, and lower in high connectivity patches in riverine aquatic landscapes relative to terrestrial landscapes. Finally at high dispersal (*d* = 1; Fig. 3 F), since virulence evolves to similar values in both landscape types, assuming similar virulence, one can predict accurately spatial distribution.

These results illustrate how knowing evolutionary optima for parasite virulence can allow us to make general predictions about differences in spatial parasite distribution across landscape types. Thus overall, by accounting for evolutionary stable virulence, we show that we should find greater local parasite extinction in riverine aquatic landscapes (Fig. S14). Interestingly, since most of these extinctions are located in degree-1 patches (Fig. 3 D–F), this decreases the likelihood that the parasite goes extinct globally, relative to terrestrial landscapes in which local extinctions are more are less variable across connectivity. Other examples, namely parasite prevalence and host population densities (Fig. S15) show that differences in virulence evolution between the two landscape types do not change predictions relative to what is expected for a fixed virulence.

Thus, with these examples, we highlight the complex interplay between landscape structure, evolutionarily stable virulence and predicting epidemiological variables.

## General discussion

In summary, we find that virulence evolves differently in terrestrial and riverine aquatic landscapes, with kin selection as the main evolutionary driver. Further, by analysing neutral vatiation we show that the two landscapes produce characteristic patterns of parasite relatedness that explain differences in virulence evolution. We further show how network topology, through the specific patterns of patch connectivity and their impact on demography, generates differences in parasite relatedness between terrestrial and aquatic landscapes. Ultimately, the interplay between network-specific ecological effects and network-specific evolutionarily stable virulence defines the spatial distribution of hosts and parasites, and more broadly host-parasite metapopulation dynamics Our modelling exercise allows us to identify a mechanistic eco-evolutionary feedback loop (Fig. 4; Govaert et al. 2019; Fronhofer et al. 2023). Ecologically, landscape structure impacts dispersal patterns of individual hosts and their parasites, leading collectively to characteristic host-parasite spatial dynamics. These spatial network specific dynamics further define patterns of parasite relatedness in the landscape, which impacts the evolution of parasite virulence (eco-to-evo). Parasite virulence evolution in turn impacts the demographic rates of the hosts and parasites driving ecological patterns at the level of the patch (e.g., patterns of local parasite extinction), but also the distribution of hosts and parasites at the landscape scale (evo-to-eco).

**Figure 4:**
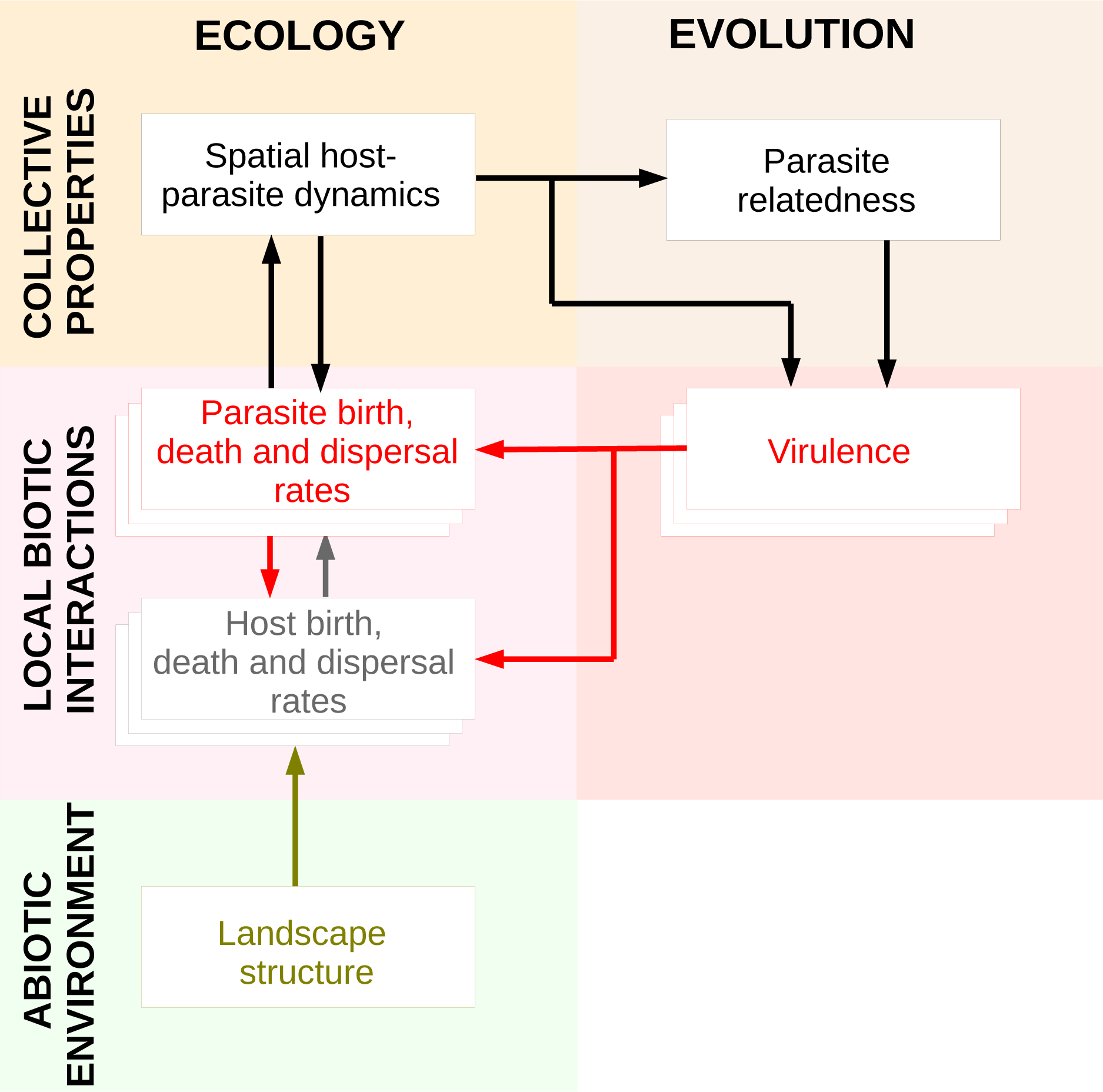
An eco-evolutionary feedback between host-parasite demography, landscape structure and virulence evolution. Clearly, landscape structure represents the abiotic environment, the biotic host-parasite interactions are shown by stacked boxes of hosts and parasites, ecological interactions and evolutionary dynamics represented by the virulence genotypes. Together, the abiotic environment, demography and evolution lead to collective properties at a higher level of organisation that can further feed back onto demographic rates. In our study, the landscape context is the abiotic environment that impacts host and parasite dispersal, since it impacts host (hence, parasite) dispersal patterns, and also their demography by modifying their spatial distribution. Virulence is genetically encoded, and it directly impacts parasite demography, via the virulence-transmission trade-off, and host demography by reducing the number of offspring the host produces (evo-to-eco). Finally, patterns of host-availability and parasite relatedness that result from host-parasite demography feed back onto parasite demographic rates, which, via selection and gene flow, impact trait evolution (eco-to-evo).

More generally, this feedback loop allows us to derive the empirically testable prediction that parasite virulence as a function of host dispersal should differ generally between terrestrial and aquatic landscapes (Fig. 1). This prediction remains untested despite recent efforts to quantify impacts of parasites on host reproduction (Hasik and Siepielski, 2022) in a meta-analysis since data are mostly available on terrestrial systems. Of course, natural host-parasite systems likely also have other drivers of virulence than landscape structure. Additionally, riverine aquatic and terrestrial parasites could differ generally in their life cycles. While we recognise these limitations, our model shows that, all else being equal, the landscape context alone can modify eco-evolutionary outcomes. Below, following the framework developed in Fig. 4, we will discuss our work in the context of previous studies on virulence evolution, ecological dynamics in complex networks and evo-to-eco feedbacks in host-parasite systems.

### Eco-to-evo: Parasite relatedness, hence, strength of kin selection results from network topology

Focusing the eco-to-evo component of Fig. 4, we emphasise the role of kin selection for virulence evolution in spatially structured systems. It has previously been shown in infinite island models (Wild et al., 2009) and using pair approximations in lattice models (Lion and Boots, 2010) that virulence evolution in spatially structured systems should be driven by kin selection. In Wild et al. (2009), increasing parasite dispersal leads to greater virulence due to reduced competition with kin locally (kin shading). Thus, evolution of reduced virulence can be thought of as an altruistic strategy (Lion and Boots, 2010). While these studies assume virulence acting on host mortality which behaves differently from our assumption of fecundity virulence (Abbate et al., 2015), how spatial structure impacts both these kinds of virulence is similar (O’Keefe and Antonovics, 2002). Generally, the simplifying assumptions on spatial structure allow for an analytical expression for selection gradients to be obtained. Thus, the mechanisms driving evolution of parasite virulence going from well mixed to simplified systems are well worked out. While modelling more complex spatial structure does not allow for analytical treatment, it is possible to partition individual and inclusive fitness components by re-shuffling simulations (Poethke et al., 2007; Deshpande et al., 2021), like in the present study, confirming the role of kin selection. Placed within this context, we show that parasite relatedness, hence the strength of kin selection itself results from spatial network topology, which determine local parasite relatedness, further driving the evolution of parasite virulence.

While it is now recognised that terrestrial and aquatic landscapes have different properties (Rinaldo et al., 2014; Carraro et al., 2020; Gilarranz, 2020), we show that these properties lead to different evolutionary dynamics. Previous work has investigated the impact of modularity, which is a feature of terrestrial landscapes, and how it slows the spread of spatial perturbations (Gilarranz et al., 2017). Studies on riverine networks have focused theoretically and experimentally on metapopulation (Fronhofer and Altermatt, 2017) and metacommunity (Carrara et al., 2012) dynamics. Abstract representations of riverine landsapes have also been used in a more applied perspective to show differences in parasite spread upstream and downstream such landscapes (Carraro et al., 2018). While these studies focus only on one landscape type, Saade et al. (2023) have recently studied bistabilty and hysteresis comparing terrestrial and aquatic metapopulations. Going beyond this, we show that not only should ecological dynamics differ between terrestrial and riverine aquatic landscapes, but also patterns of selection and gene flow, leading to differences in trait evolution. Therefore, the patterns of kin selection reported here are only one example of how landscape structure impacts eco-evolutionary feedbacks. Specifically in our study, by generating an expectation of parasite relatedness that depends on landscape structure and dispersal, but not on evolved virulence, we can investigate the impact of ecology (landscape structure) on patterns of parasite relatedness hence, virulence evolution by breaking the feedback of evolution on ecology.

Our result that it is the heterogeneity in connectivity of riverine networks that leads to characteristic patterns of parasite relatedness can be placed in context of previous work on patterns of neutral genetic and species diversity in spatial networks. Boussange and Pellissier (2022) study this effect of network structure on neutral genetic diversity and its consequences on evolution of local adaptation to an environment. They show that neutral genetic differentiation in spatial networks depends on both connectivity and variation in population densities due to heterogeneous degree distributions. Similarly, it has been shown theoretically that dendritic networks maintain greater genetic diversity compared to grid landscapes (Paz-Vinas and Blanchet, 2015; Thomaz et al., 2016). Parallels can also be drawn to characteristic patterns of species diversity in spatial networks (Economo and Keitt, 2008) and particularly riverine networks (Muneepeerakul et al., 2007; Carrara et al., 2012). Finally, spatial network structure has also been shown to drive the evolution of dispersal (Muneepeerakul et al., 2011; Henriques-Silva et al., 2015; Fronhofer and Altermatt, 2017).

Importantly, while going from a simplified spatial context (such as infinite island models), to more complex spatial structures, the basic mechanisms driving trait evolution do not change. However, landscape structure acts to modulates them (Fig. 4). In a host-parasite context we here study an illustrative example of virulence evolution, however, patterns of local adaptation as a result of co-evolution also critically depend on gene flow (Gandon and Michalakis, 2002) within networks (Gibert et al., 2013; Jousimo et al., 2014; Höckerstedt et al., 2022). Beyond host-parasite systems, linking our results to kin selection also implies that landscape structure should modulate the evolution of social behaviour, in general.

### Evo-to-eco: Differences in virulence evolution may amplify, override or reduce differences between terrestrial and aquatic landscapes at different spatial scales

Finally, taking into account how virulence evolution differs between terrestrial and riverine aquatic landscapes, we can predict the spatial distribution of hosts and parasites. As discussed above, even without differences in virulence evolution, terrestrial and aquatic landscapes are generally expected to differ in their ecological properties (see for example, Saade et al. 2023). Fig. 3 re-emphasises this point, where the fixed virulence scenario shows differences in the spatial distribution of local parasite extinction between terrestrial and aquatic landscapes. First, this shows that ecological variables as a function of patch degree can only be understood in the context of the entire landscape, since for the same connectivity they differ between two landscape types. Second, taking an evolutionary perspective allows us to understand how differences in trait evolution can increase, decrease or not impact ecological variables given similar connectivity in different landscapes. Thus, accurate predictions of disease distributions and spread may require information on landscape topology. Our work demonstrates how virulence evolution can in turn impact host-parasite ecological dynamics (e.g. patterns of local parasite extinction) at different spatial scales (evo-to-eco). Considering virulence evolution in terrestrial and aquatic landscape allows us to show how differences in the spatial distribution of hosts and parasites between such landscapes can be reduced (in the case of local parasite extinction, and intermediate dispersal), amplified (in the case of local parasite extinction, and low dispersal) or stay the same (host population density and parasite prevalence for all dispersal rates), as a consequence of virulence evolution. This complex interaction can be explained based on Fig. 4, since landscape structure directly impacts the spatial distribution of hosts and parasites. Because this in turn impacts how virulence evolves, the consequences of virulence evolution are indirect, and differences in landscape structure alone may dominate. At the level of the metapopulation, landscape structure clearly dominates over differences in evolved virulence for parasite persistence (global parasite extinction), with global extinctions being greater for terrestrial landscapes even though evolved virulence is lower. Thus, the interaction between landscape type and evolution must be considered when trying to predict spatial epidemiology. Recent work by Jousimo et al. (2014) and Höckerstedt et al. (2022) support this finding, since the authors show that increased host-gene flow in highly connectivity patches leads to increased resistance, reducing harm to the host, and lower parasite occupancy.

### Future direction

While our study highlights how taking into account spatial network structure impacts our understanding of eco-evolutionary feedbacks in host-parasite systems, some questions remain. For example, in our study we assumed that dispersal is fixed and cannot evolve, implying that our model applies to host-parasite systems in which the parasite evolves much faster than the host as is likely in host parasite systems where number of parasites is much greater than the host. However, host dispersal may also evolve in other kinds of systems and more importantly can be driven by landscape structure (Muneepeerakul et al., 2011; Henriques-Silva et al., 2015; Fronhofer and Altermatt, 2017) which can be explored in detail in further work. We also do not account for riverine directionality as this makes ecological and evolutionary patterns even stronger (Fronhofer and Altermatt, 2017), but a detailed investigation of more realistic riverine landscapes is possible.

## Conclusion

In conclusion, by studying how parasite virulence evolves in terrestrial and riverine aquatic landscapes, we highlight an eco-evolutionary feedback between landscape structure, host-parasite demography and parasite virulence (Fig. 4). It is now increasingly recognised that eco-evolutionary dynamics of host-parasite systems (Rogalski et al., 2017) are greatly modified by anthropogenic change leading to phenomena such as emerging infectious diseases (Daszak et al., 2000). Particularly, landscapes are vulnerable to changes such as habitat fragmentation and increased barriers to dispersal, generally, rewiring of dispersal networks (Bullock et al., 2018). Such disturbances maybe intrinsically (due to landscape structure) or extrinsically (due to the nature of human activities), differ between terrestrial and riverine aquatic systems (Johnson and Paull, 2011), and thus might require differing management effort. Our work highlights that an eco-evolutionary perspective may be needed for a complete understanding and an efficient management of such systems.

## Acknowledgements

We thank Śebastien Lion, Fraņcois Massol and Fraņcois Rousset for helpful discussion of the results. This work was funded by a grant from the Agence Nationale de Recherche to O.K. (grant no. ANR-20-CE02-0023-01). E.A.F has received financial support from the CNRS through the MITI interdisciplinary program. This is publication ISEM-YYYY-XXX of the Institut des Sciences de l’Evolution – Montpellier

