## Supplementary Material for "Landscape structure drives eco-evolution in host-parasite systems"

### Supplementary methods

### Model description

#### Model overview

We develop a spatially explicit individual-based SI (Susceptible-Infected) model of an asexual host with discrete non-overlapping generations infected by an asexually reproducing parasite. We only explicitly model the host species, but introduce features described below to our model such that we can still capture parasite evolutionary dynamics. Thus, henceforward, the word "individual" only refers to the host species. An individual is said to be susceptible if it does not bear a parasite and it is said to be infected if it does. Parasite virulence ($v$) is a haploid trait that evolves as a by-product of increasing transmission. Thus, we assume an increasing relationship between virulence and transmission. Virulence in our model acts by reducing the fecundity of the host (Abbate et al., 2015), if it is infected. At the end of each generation, the infected hosts release parasite propagules that can infect susceptible hosts from the offspring generation. Thus, transmission is horizontal. We model landscapes as a spatial network of $n = 100$ discrete patches, each with local host density regulation and parasite transmission. The connectivity between patches is given by the connectivity matrix $\mathbf{D}_{n\times n}$ and this matrix varies for the different landscape types explored. Individuals may disperse between patches with a fixed probability $d$. Our primary analysis focuses on differences between virulence evolution in terrestrial (represented by random-geometric graph; RGG) and aquatic riverine (represented by optimal channel networks; OCNs) landscapes.

#### Life cycle

##### Dispersal

After they are born, host individuals first disperse with a fixed dispersal probability $d$ to one of their neighbours with equal probability. Whether an individual can disperse from its natal patch $x$ to another patch $y$ is given by an undirected unweighted landscape connectivity matrix $\mathbf{D}_{n\times n} = (D_{xy})$. $D_{xy} = 1$ if individuals can disperse from patch $x$ to patch $y$, and if they cannot then $D_{xy} = 0$.

##### Host reproduction

After dispersal, individuals reproduce in their target patches. The number of offspring each individual produces is given by the Beverton-Holt model of logistic growth (Beverton and Holt, 1957). After emigration and immigration, if there are $S_x(t)$ susceptible, $I_x(t)$ infected individuals and $N_x(t) = S_x(t) + I_x(t)$

Table S1: Model Parameters/Variables. Model parameters for the focal scenario are in bold.

| Model Parameter/Variable | Description | Values |
|---|---|---|
| $N_x(t)$ | Total population size of host in patch $x$ at time $t$ | Dynamic |
| $S_x(t)$ | Total population size of susceptible hosts in patch $x$ at time $t$ | Dynamic |
| $I_x(t)$ | Total population size of infected hosts in patch $x$ at time $t$ | Dynamic |
| $\lambda_0$ | Intrinsic population growth rate of host in Beverton-Holt model | $2, \mathbf{4}$ |
| $\alpha$ | Intra-specific competition coefficient of host in Beverton-Holt model | $\mathbf{0.01}$ |
| $d$ | Dispersal probability of host | $\mathbf{0.01}, \mathbf{0.05}, \mathbf{0.1}, \mathbf{0.2}, \mathbf{0.3}, \mathbf{0.4}, \mathbf{0.5}, \mathbf{0.6}, \mathbf{0.7}, \mathbf{0.8}, \mathbf{0.9}, \mathbf{1}$ |
| $v$ | Parasite virulence as reduced host fecundity | Evolves during simulations and initialised with standing genetic variation |
| $\beta(v)$ | Parasite transmission rate | Evolves during simulations |
| $\beta_{max}$ | Maximum possible parasite transmission rate | $4, \mathbf{8}, 10$ |
| $s$ | Shape of the relationship between parasite virulence and transmission | $\mathbf{0.5}, 1, 2$ |
| $a$ | Searching efficiency of the parasite | $\mathbf{0.01}$ |
| $m_v$ | Mutation rate of parasite virulence | $\mathbf{0.01}$ |
| $\sigma_v$ | Effect size (standard deviation) of mutation after logit transform | $\mathbf{0.5}$ |

individuals in a given patch $x$ in the generation $t$, then in the next generation $t+1$, the expected number of offspring is given by:

$$N_x(t+1) = \frac{\lambda_0 S_x(t)}{1+\alpha N_x(t)} + \frac{(1-v)\lambda_0 I_x(t)}{1+\alpha N_x(t)}. \tag{S1}$$

Here, $\lambda_0$ is the intrinsic population growth rate, $\alpha$ is the intra-specific competition coefficient, $v$ is the expected virulence in the patch defined here as reduction in fecundity of an infected host. The realised number of offspring for each individual $k$ in a patch $x$ is drawn from a Poisson distribution with $\frac{\lambda_0}{1+\alpha N_x(t)}$ if susceptible and $\frac{(1-v_k)\lambda_0}{1+\alpha N_x(t)}$ if infected, where $v_k$ is the virulence of the parasite that infects the given individual in the patch.

#### Parasite transmission

The parental generation at generation $t$ dies after reproduction. The dead infected individuals from the parental generation in a given patch release parasite propagules which can then infect new-born individuals of the offspring generation at generation $t+1$. The expected number of infected individuals $I_x(t+1)$ at generation $t+1$ in a patch $x$ is given by a modified Nicholson-Bailey model (Nicholson and Bailey, 1935) with a Holling type II functional response (Chaianunporn and Hovestadt, 2012; Deshpande et al., 2021):

$$I_x(t+1) = N_x(t+1)\left(1 - \exp\frac{-a\beta(v)I_x(t)}{1 + aN_x(t+1)}\right). \tag{S2}$$

Here, $\beta(v)$ is the expected number of parasite propagules released by an infected individual and $a$ is the searching efficiency of these propagules. Similar to previous work (Boots and Sasaki, 1999; O'Keefe and Antonovics, 2002; Lion and Boots, 2010) we assume an increasing relationship between virulence and transmission. This assumption is consistent with the trade-off hypothesis (reviewed in Alizon et al., 2009). Therefore, we can write $\beta(v)$ as:

$$\beta(v) = \beta_{max}v^s. \tag{S3}$$

Here, $\beta_{max}$ represents the maximum possible transmission rate and $s$ determines how rapidly transmission increases with virulence, that is, the shape of the trade-off curve. Saturating, linear and accelerating curves are represented by $s < 1$, $s = 1$ and $s > 1$ respectively. The expected number of susceptible individuals in the $t+1$ generation is therefore given by $S_x(t+1) = N_x(t+1) - I_x(t+1)$. We assume that whether a given parasite propagule released by a parental individual $k$ comes in contact with a newborn susceptible offspring is Bernoulli distributed with probability $1 - \exp\frac{-a\beta(v_k)}{1+aN_x(t+1)}$. This means that multiple parasite propagules can come in contact with a single newborn host. We assume that if a newborn host is in contact with multiple parasite propagules, it can ultimately be infected only by one such propagule which is drawn with equal probability.

The parasite mutates within the newborn host, thus we add a mutation with a probability $m_v$. Since the virulence trait can only take values between 0 and 1, we perform a logit transformation of the mutated virulence to ensure that the virulence trait remains constrained. Thus, mutation effect for logit transformed virulence is drawn from a normal distribution with standard deviation $\sigma_v$. An inverse logit transformation is then performed on the mutated virulence genotype to get realised mutation effect. In the beginning of the simulation all the haploid virulence loci are initialised with values drawn from a uniform distribution on the interval $[0, 1]$, representing standing genetic variation. Each parasite also contains a haploid neutral marker locus which is transmitted during infection, with the same mutation rate and effect as the virulence trait. The neutral marker locus is also initialised similar to virulence and with the same mutation rate. Adding this locus allows us to track patterns of relatedness between parasites within the metapopulation (described in a later section).

#### The landscapes

##### Terrestrial landscapes and controls

Terrestrial landscapes are represented by random-geometric graphs (RGGs). Generally, we assume that patches are uniformly distributed in space and dispersal is possible only between patches within a given radius. These assumptions lead to modularity in RGGs (Gilarranz, 2020; Saade et al., 2023). Thus, RGGs are generated by drawing $n$ points on a unit square $[0,1]^2$ from a uniform distribution and joining them if they are closer than a specified distance. We use the R package igraph (version 1.2.11) to generate these landscapes.

We generate an ensemble of 1000 realisations with radius $r = 0.15$ implying a realised average degree 6.08 with the condition that networks are connected. The average degree is defined as the average number of patches to which a patch in a network is connected. To tease apart the effect of average degree alone vs. network topology, we design landscapes with same average degree. RGGs have the following properties: they are spatially structured, have heterogeneity in patch degree and are modular (Gilarranz, 2020). The controls therefore are: 1) random networks (average degree 5.98; not spatially structured or modular but they have heterogeneity in patch degree), 2) hexagonal regular grids (average degree 6; which are spatially structured but have homogeneous degree distribution and lack a modular structure) and maximally 3) maximally modular networks (they are spatially structured and modular but have mostly homogeneous degree distribution). Thus, differences in virulence evolution between RGGs and control landscapes imply that average degree alone does not explain virulence evolution in RGGs.

##### Riverine aquatic landscapes and controls

Here, we focus on riverine landscapes that are represented by optimal channel networks (OCNs). OCNs take into account geomorphological processes (Rinaldo et al., 2014) that are thought to shape river networks globally. We generate river structures organised on a $10 \times 10$ grid using the R package OCNet (version 0.5.0; Carraro et al. 2020). We fix the position of the outlet to an outer patch in the grid, and we do not aggregate the patches or take into account the direction of flow. Since more upstream patches are also likely to have a lower connectivity, directional flow upstream to downstream only tends to amplify patterns that are obtained in the absence of directional flow. OCNs have previously been used to study metapopulation dynamics (Fronhofer and Altermatt, 2017) and disease spread (Carraro et al., 2018) in riverine systems. OCNs have a dendritic structure, hence, a highly heterogeneous degree distribution, however the average degree of these landscapes is 1.98 (Saade et al., 2023). Thus, to pin down the effect of average degree alone, we design control landscapes that have an average degree close to riverine aquatic landscapes but differing degree distribution. Particularly, compare virulence evolution in riverine aquatic landscapes to: 1) circular landscapes in which each patch has exactly 2 neighbours and 2) a "spiky" network generated by adding 68 patches of degree 1 to an Erdös-Renyi network of average degree 4.13 with 32 patches (realised average degree 2.67). These two networks have different topology but have heterogeneous degree distribution with many degree 1 patches (68.45 in OCN and 68.20 in the "spiky" landscape).

#### Model analysis

We extract the median evolved parasite virulence at the end of 2500 time steps which is sufficient for evolutionary dynamics to reach (quasi)-equilibrium (Fig. S1). We run our model for 1000 replicate simulations in our focal scenario, for varying host dispersal probability in terrestrial (RGG) and aquatic landscapes (OCNs) along with the control landscapes (circular, spiky, grid, random and modular controls). We run simulations for all parameter combinations in Table S1, but we present results for one focal scenario, highlighted in bold in the same table.

#### Testing the role of kin selection

##### Re-shuffling of parasites in landscapes

We run additional simulations in which, for each time step, after parasite transmission has taken place, we collect all parasite genotypes (virulence and neutral locus values associated with an infected individual) in the landscape and re-distribute these parasites randomly across the landscape, while maintaining the same number of total and infected densities per-patch. This breaks kin structure while keeping epidemiological structuring the same. This simulation experiment has previously been used by Poethke et al. (2007) and Deshpande et al. (2021) to separate out individual vs. inclusive fitness components.

##### Measuring parasite relatedness

To understand the mechanism for virulence evolution, since virulence can both impact and driven by parasite relatedness and host availability (O'Keefe and Antonovics, 2002; Lion and Boots, 2010), we run additional ecological simulations, in which virulence is fixed going from $v = 0.8, 0.82, ..., 1$. For all landscapes and controls in our focal scenario and in ecological simulations, we calculate at equilibrium (last 100) time-steps the average parasite relatedness within a patch. We also calculate the geometric mean over the last 100 time steps of host densities within a patch. Thus, we can we can quantify the long term changes in host availability within a patch. We calculate the relatedness between two randomly chosen parasite strains in patch $x$ and $y$ as follows:

$$R = \sum_k p_{k,x} q_{k,y} \tag{S4}$$

Where $p_{k,x}$ is the frequency of a strain $k$ of the neutral locus in patch $x$ and $q_{k,y}$ its frequency in patch $y$. this gives us the average probability that two strains chosen at random with replacement in patches $x$ and $y$ in a patch are identical by state. So, average relatedness within a patch in a metapopulation of $n$ patch is given by $R_{home} = \frac{1}{n} \sum_x \sum_k p_{k,x}^2$

### Supplementary figures

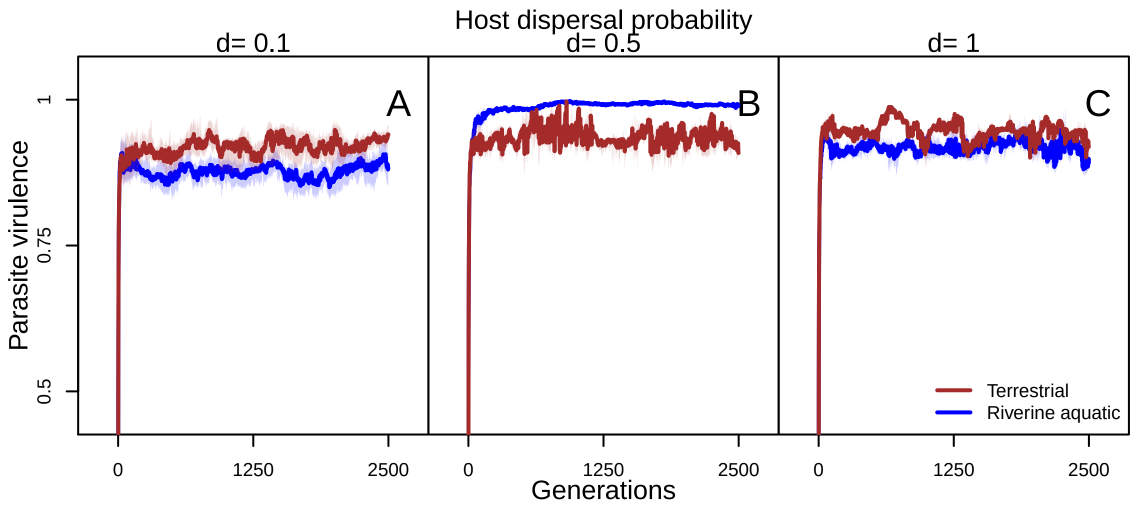


Figure S1: Example dynamics of virulence evolution in terrestrial (brown) and aquatic (blue) landscapes for different host dispersal probability ($d = 0.1, 0.5, 1$). We plot the measured parasite virulence over time for one example replicate simulation. Solid lines represent the median virulence for a generation over all patches in the landscape, and the shaded region represents interquartile range. The evolutionary dynamics attain (quasi)-equilibrium in both terrestrial and riverine aquatic landscapes. Model parameters of the focal scenario: intrinsic growth rate of the host $\lambda_0 = 4$, intraspecific competition coefficient of the host $\alpha = 0.01$, maximum parasite transmission $\beta_{max} = 8$, shape of trade-off curve $s = 0.5$, searching efficiency of parasite $a = 0.01$.

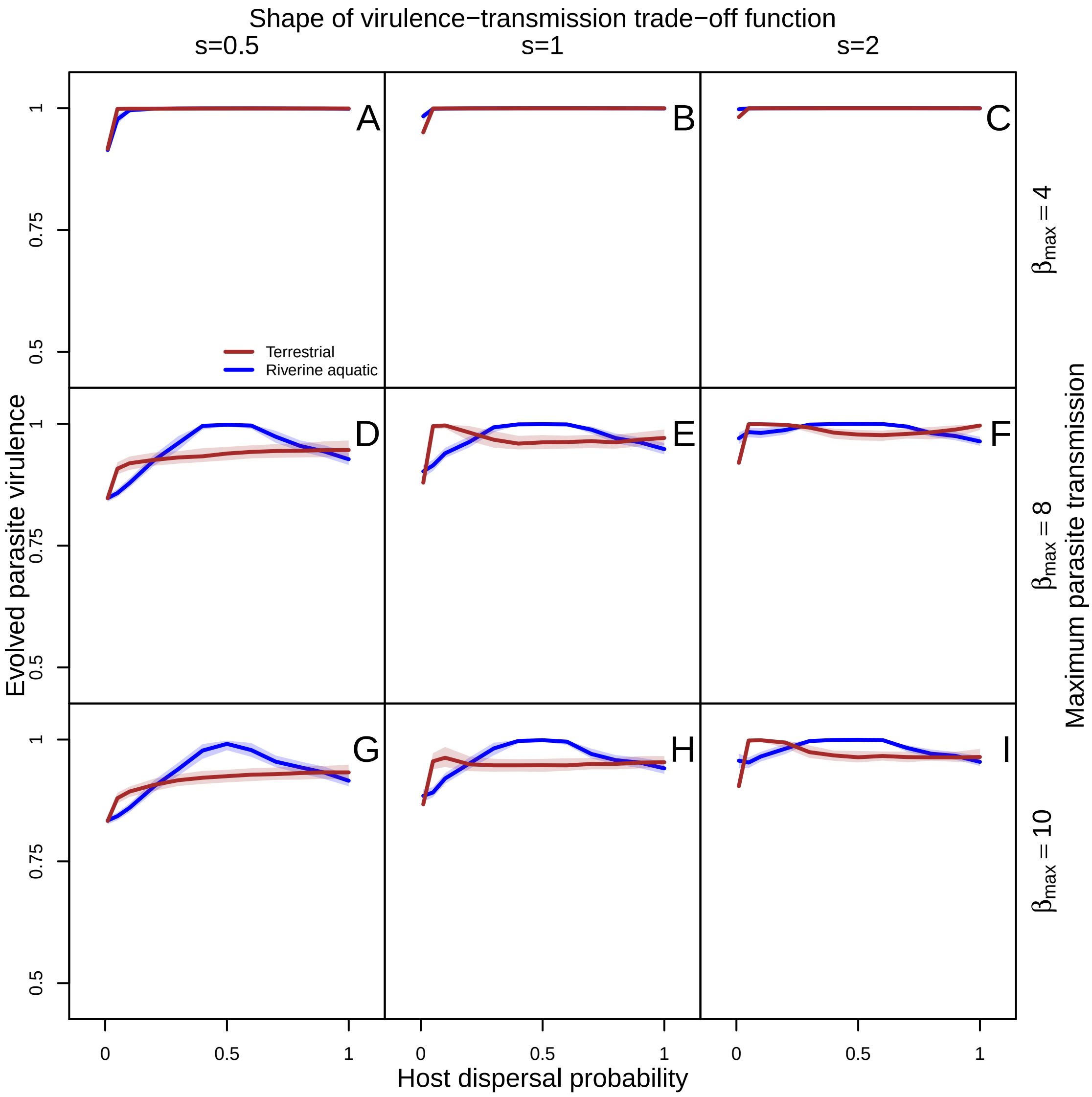


Figure S2: Sensitivity to maximum transmission ($\beta_{max}$) and shape of virulence-transmission trade-off function of evolved parasite virulence as a function of host dispersal ($d = 0.01, 0.05, 0.1, 0.2, 0.3, 0.4, 0.5, 0.6, 0.7, 0.8, 0.9, 1$) in terrestrial (brown) and riverine aquatic (blue) landscapes. From left to right: shape of the the virulence transmission trade-off changes $s = 0.5, 1, 2$ implying that transmission is a saturating, linear and accelerating function respectively of virulence. From top to down, the maximum possible transmission rate increases ($\beta_{max} = 4, 8, 10$). The solid line represents the median and shaded regions the inter-quartile range of the evolved virulence trait over 1000 replicate simulations, median over all parasites at the last simulation time step ($t = 2500$). The basic pattern that virulence as a function of host dispersal depends on landscape structure holds for larger $\beta_{max} = 8, 10$ for all shapes of the virulence transmission trade-off function. Model parameters: intrinsic growth rate of the host $\lambda_0 = 4$, intraspecific competition coefficient of the host $\alpha = 0.01$, searching efficiency of parasite $a = 0.01$.

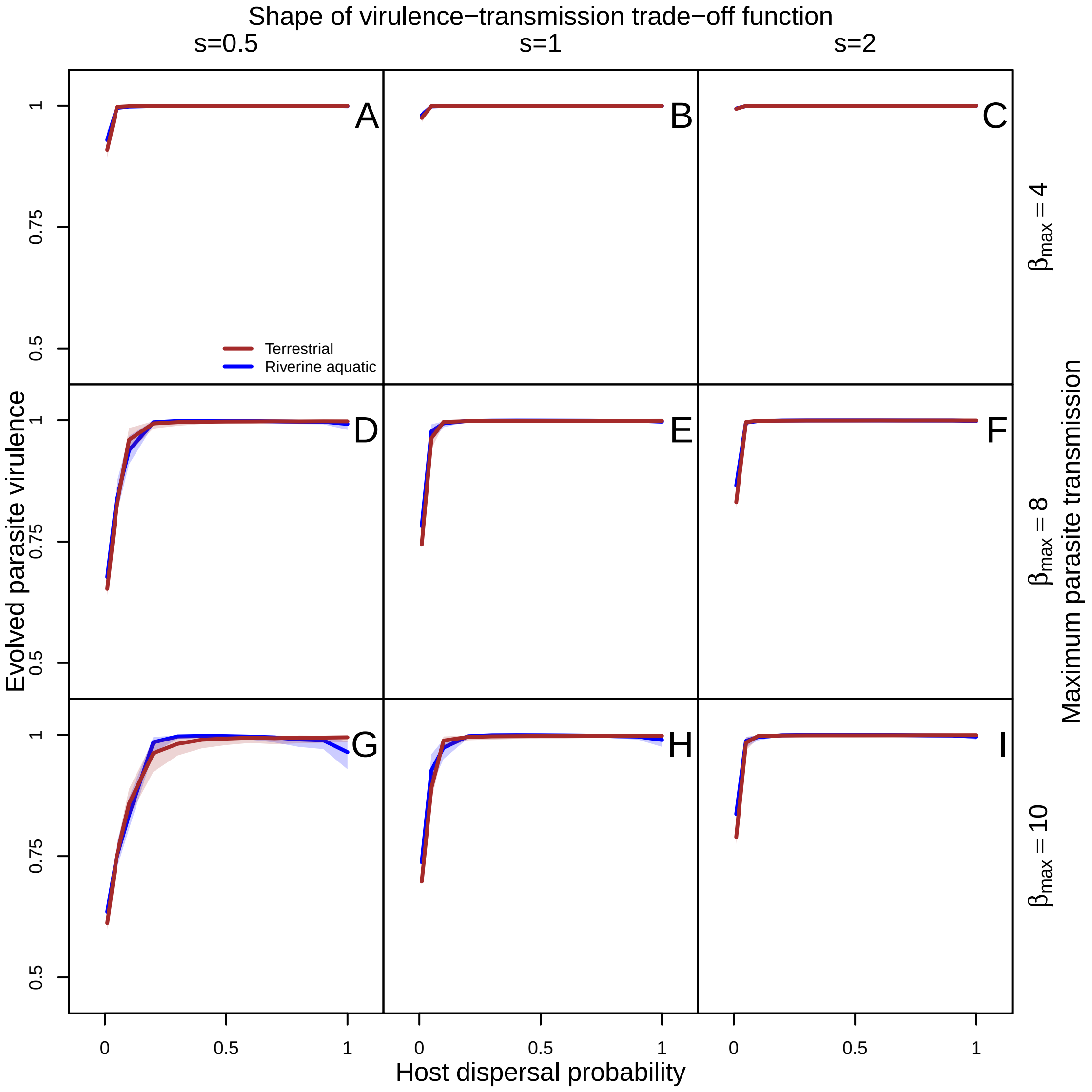


Figure S3: Sensitivity to lower host intrinsic growth rate ($\lambda_0 = 2$) of evolved parasite virulence as a function of host dispersal ($d = 0.01, 0.05, 0.1, 0.2, 0.3, 0.4, 0.5, 0.6, 0.7, 0.8, 0.9, 1$) in terrestrial (brown) and riverine aquatic (blue) landscapes. From left to right: shape of the the virulence transmission trade-off changes $s = 0.5, 1, 2$ implying that transmission is a saturating, linear and accelerating function respectively of virulence). From top to down, the maximum possible transmission rate increases ($\beta_{max} = 4, 8, 10$). The solid line represents the median and shaded regions the inter-quartile range of the evolved virulence trait over 1000 replicate simulations, median over all parasites at the last simulation time step ($t = 2500$). At lower host growth rate $\lambda_0 = 2$, virulence primarily depends on host dispersal alone and not on landscape structure. Model parameters: intrinsic growth rate of the host $\lambda_0 = 2$, intraspecific competition coefficient of the host $\alpha = 0.01$, searching efficiency of parasite $a = 0.01$.

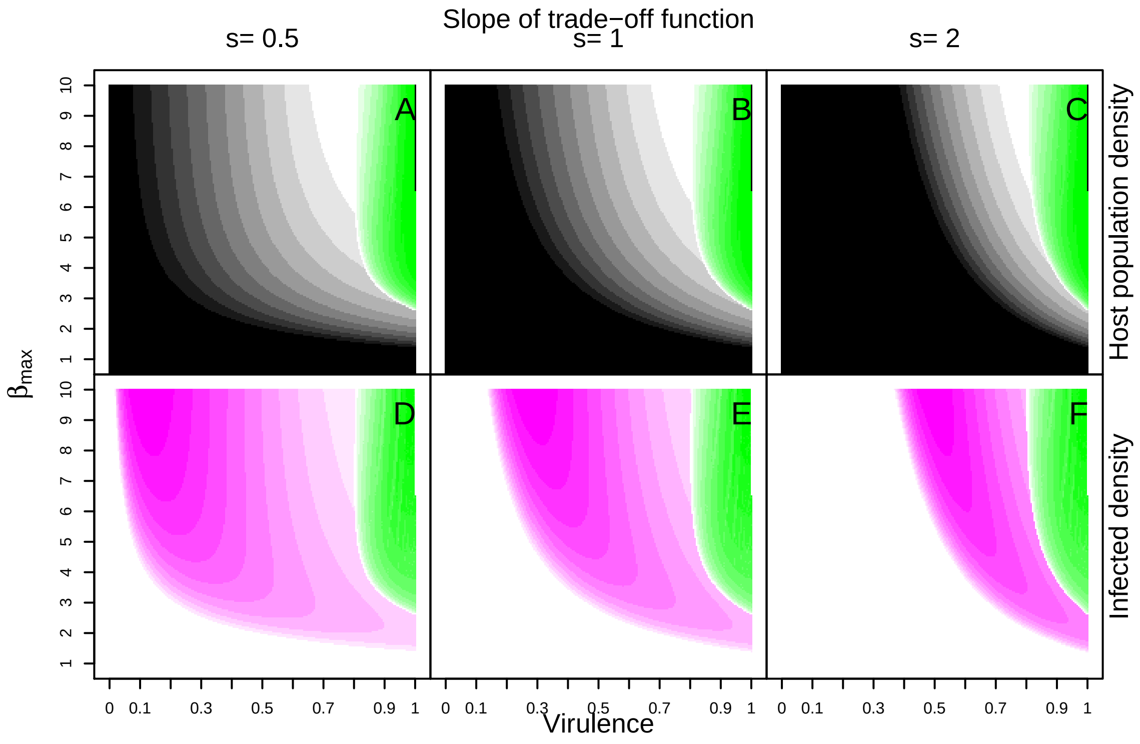


Figure S4: Deterministic one patch dynamics of the host-parasite model. From left to right: shape of the the virulence transmission trade-off changes $s = 0.5, 1, 2$ implying that transmission is a saturating, linear and accelerating function respectively of virulence). A–C: Steady state host population density plotted as a function of fixed virulence and maximum transmission $\beta_{max}$ parameters. Darker shades of black indicate larger host population densities. The green region represents the area of parameter space where oscillations occur in host population densities, with darker shades indicating a higher amplitude as represented by the difference between that maximum and minimum host densities. D–F: Steady state infected density plotted as a function of fixed virulence and maximum transmission $\beta_{max}$ parameters. Darker shades of pink indicate larger host population densities. The green region represents the area of parameter space where oscillations occur in infected densities, with darker shades indicating a higher amplitude as represented by the difference between that maximum and minimum infected densities. In the absence of spatial structure, virulence is expected to evolve to a maximal value of $v = 1$ (O'Keefe and Antonovics, 2002), which is in the oscillatory regime. In terrestrial and aquatic landscapes all evolved virulences also fall in this regime. However, since at lower transmission rates, the amplitude of oscillations is not large enough to reduce susceptible host densities except at very low dispersal rates, there is little cost to parasite fitness due to increasing virulence, leading to evolution of maximal virulence. Model parameters: intrinsic growth rate of the host $\lambda_0 = 4$, intraspecific competition coefficient of the host $\alpha = 0.01$, searching efficiency of parasite $a = 0.01$

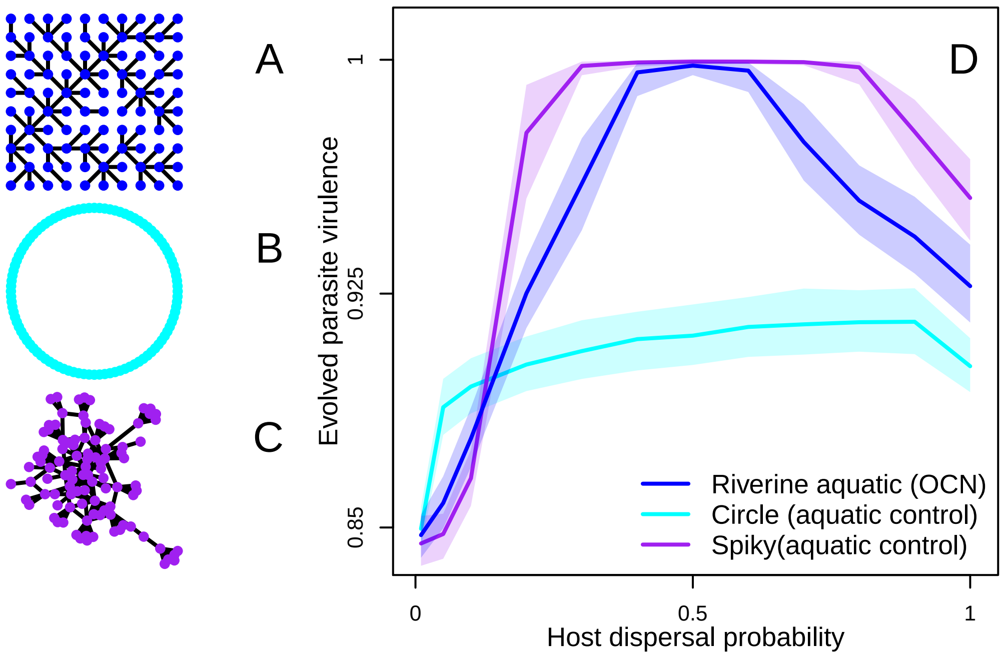


Figure S5: Disentangling the effect of average degree vs. higher order properties on virulence evolution in riverine aquatic landscapes. Evolved parasite virulence as a function of host dispersal ($d = 0.01, 0.05, 0.1, 0.2, 0.3, 0.4, 0.5, 0.6, 0.7, 0.8, 0.9, 1$) in A: riverine aquatic (OCN, blue), B: circular (light blue) and C: spiky (purple) landscapes. D: The solid line represents the median and shaded regions the inter-quartile range of the evolved virulence trait over 1000 replicate simulations, median over all parasites at the last simulation time step ($t = 2500$). The qualitative patterns that the aquatic landscape attains maximum possible virulence ($v = 1$) at intermediate dispersal rates, and the subsequent decline is re-captured by the "spiky" network (however, virulence is over-estimated by this structure), but not by the circular network, in which the virulence saturates with dispersal to a lower value, resembling qualitatively a terrestrial landscape. This shows that average degree alone does not explain the unimodal shape of virulence as function of dispersal, but rather, the presence of a large number of degree 1 patches (68.45 in OCN and 68.20 in the "spiky" landscape) could explain this pattern. Model parameters of the focal scenario: intrinsic growth rate of the host $\lambda_0 = 4$, intraspecific competition coefficient of the host $\alpha = 0.01$, maximum parasite transmission $\beta_{max} = 8$, shape of trade-off curve $s = 0.5$, searching efficiency of parasite $a = 0.01$.

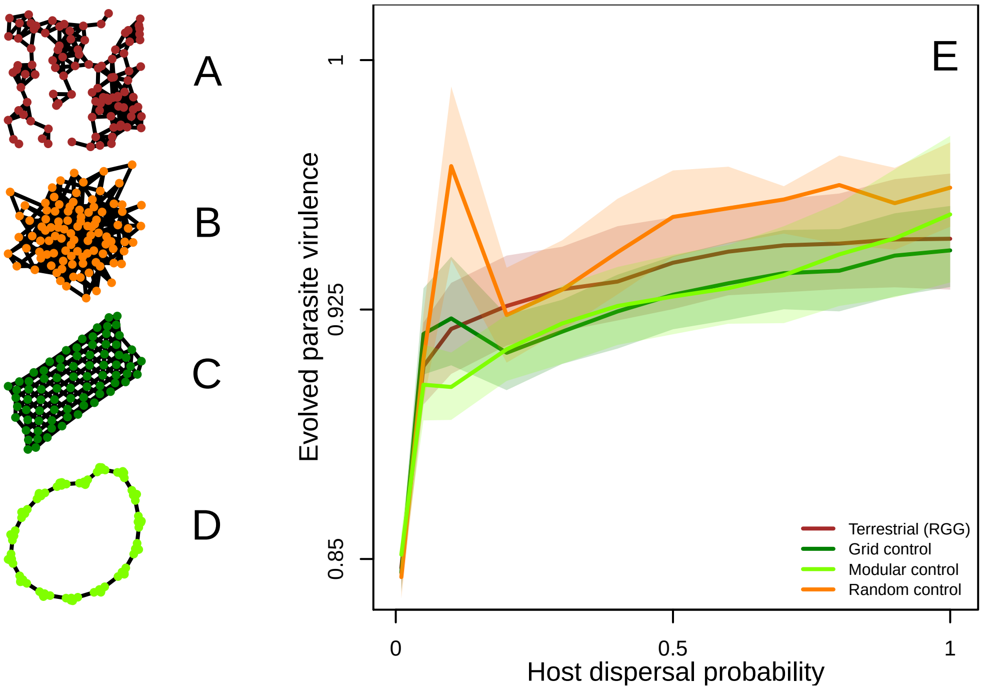


Figure S6: Disentangling the effect of average degree vs. higher order properties on virulence evolution in terrestrial landscapes. Evolved parasite virulence as a function of host dispersal ($d = 0.01, 0.05, 0.1, 0.2, 0.3, 0.4, 0.5, 0.6, 0.7, 0.8, 0.9, 1$) in A: terrestrial (RGG, brown), B: random (orange) controls, C: grid (dark green) and D: modular (light green) landscapes. E: We compare virulence evolution in terrestrial landscapes to controls with the similar average degree ($= 6.08$). The solid line represents the median and shaded regions the inter-quartile range of the evolved virulence trait over 1000 replicate simulations, median over all parasites at the last simulation time step ($t = 2500$). We find that the initial increase in virulence is greater in random networks in comparison to all other networks which are spatially structured. Model parameters of the focal scenario: intrinsic growth rate of the host $\lambda_0 = 4$, intraspecific competition coefficient of the host $\alpha = 0.01$, maximum parasite transmission $\beta_{max} = 8$, shape of trade-off curve $s = 0.5$, searching efficiency of parasite $a = 0.01$.

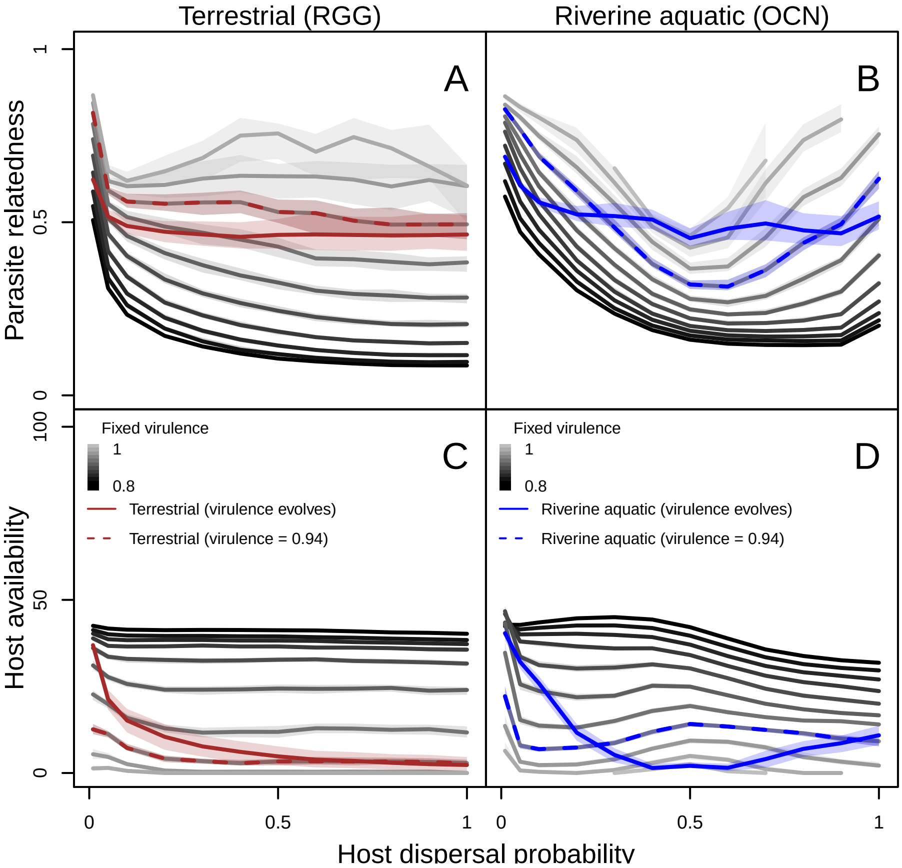


Figure S7: Parasite relatedness (A–B) and host availability (C–D) as a function of host dispersal ($d = 0.01, 0.05, 0.1, 0.2, 0.3, 0.4, 0.5, 0.6, 0.7, 0.8, 0.9, 1$) in terrestrial (RGG) (A, C) and riverine aquatic (OCN) (B, D) landscapes. Since both parasite relatedness and host availability should drive virulence evolution but also depend on it, we run additional simulations with fixed virulence ($v = 0.8, 0.82, ..., 1$; indicated by black to grey lines, the fixed virulence scenario $v = 0.94$ in the main text is indicated by dashed lines, and for each landscape type we also show relatedness and host availability when virulence can evolve) in the range of evolved virulence across all landscapes. A–B: We plot the average parasite relatedness over the last 100 time steps within a patch over all patches and (black to grey solid lines representing median over 100 replicates with increasing virulence in the ecological simulations). C–D: We plot the geometric mean of the host population density within a patch over the last 100 time steps averaged over all patches. This represents long term host availability. Model parameters of the focal scenario: intrinsic growth rate of the host $\lambda_0 = 4$, intraspecific competition coefficient of the host $\alpha = 0.01$, maximum parasite transmission $\beta_{max} = 8$, shape of trade-off curve $s = 0.5$, searching efficiency of parasite $a = 0.01$.

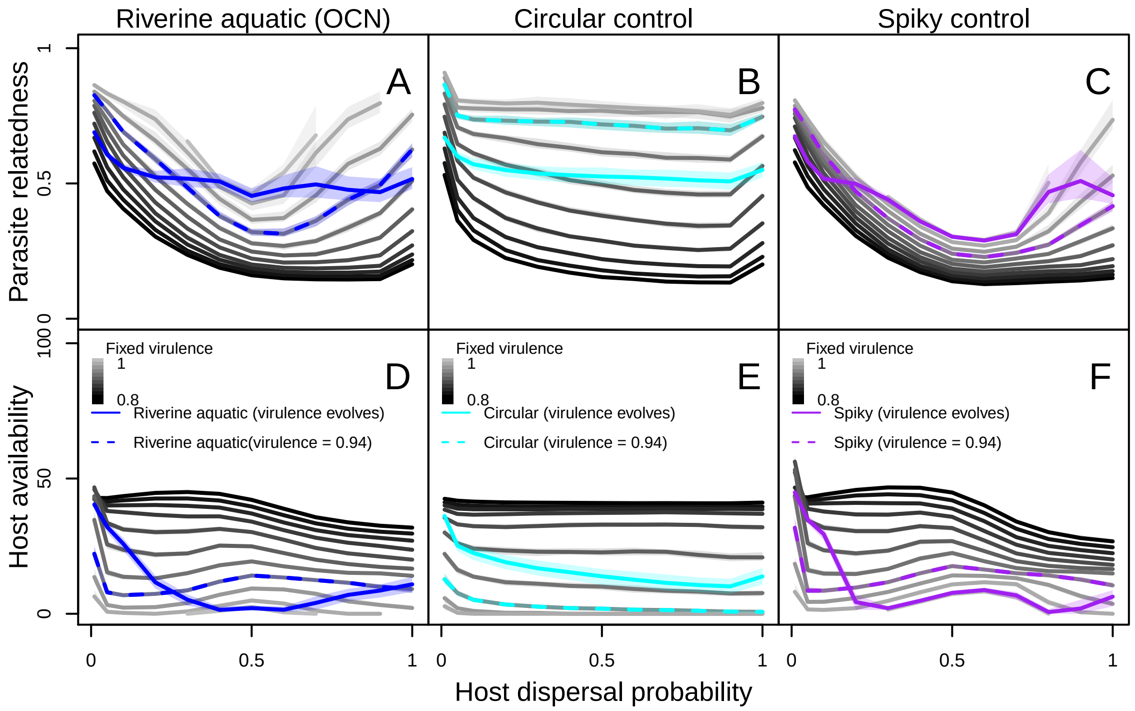


Figure S8: Parasite relatedness (A–C) and host availability (D–F) as a function of host dispersal ($d = 0.01, 0.05, 0.1, 0.2, 0.3, 0.4, 0.5, 0.6, 0.7, 0.8, 0.9, 1$) in riverine aquatic (OCN) (A, D), circular control (B, E) and "spiky" (D, F) landscapes. Since both parasite relatedness and host availability should drive virulence evolution but also depend on it, we run additional simulations with fixed virulence ($v = 0.8, 0.82, ..., 1$; indicated by black to grey lines, the fixed virulence scenario $v = 0.94$ in the main text is indicated by dashed lines, and for each landscape type we also show relatedness and host availability when virulence can evolve) in the range of evolved virulence across all landscapes. A–C: We plot the average parasite relatedness over the last 100 time steps within a patch over all patches and (black to grey solid lines representing median over 100 replicates with increasing virulence in the ecological simulations) C–D: We plot the geometric mean of the host population density over the last 100 time steps within a patch averaged over all patches. This represents long term host availability. This figure shows that patterns of parasite relatedness are consistent with evolved virulence. Particularly, relatedness is U-shaped and reaches much lower values (and virulence is unimodal and reaches much higher values) in riverine aquatic and "spiky" landscapes when compared to circular controls. Model parameters of the focal scenario: intrinsic growth rate of the host $\lambda_0 = 4$, intraspecific competition coefficient of the host $\alpha = 0.01$, maximum parasite transmission $\beta_{max} = 8$, shape of trade-off curve $s = 0.5$, searching efficiency of parasite $a = 0.01$.

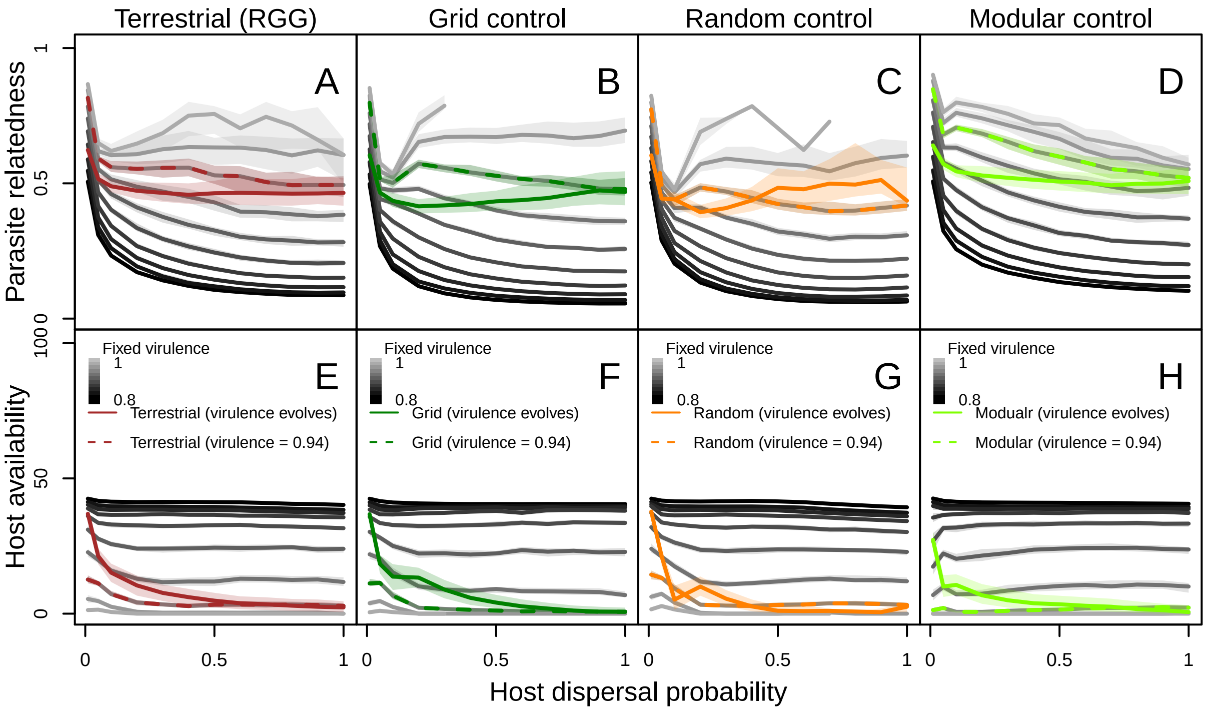


Figure S9: Parasite relatedness (A–D) and host availability (E–H) as a function of host dispersal ($d = 0.01, 0.05, 0.1, 0.2, 0.3, 0.4, 0.5, 0.6, 0.7, 0.8, 0.9, 1$) in terrestrial landscapes (RGG; A, E), grid (B, F), random (C, G) and modular (D, H) controls. Since both parasite relatedness and host availability should drive virulence evolution but also depend on it, we run additional simulations with fixed virulence ($v = 0.8, 0.82, ..., 1$; indicated by black to grey lines, the fixed virulence scenario $v = 0.94$ in the main text is indicated by dashed lines, and for each landscape type we also show relatedness and host availability when virulence can evolve) in the range of evolved virulence in the terrestrial and aquatic landscapes. A–B: We plot the average parasite relatedness over the last 100 time steps within a patch over all patches and (black to grey solid lines representing median over 100 replicates with increasing virulence in the ecological simulations) C–D: We plot the geometric mean of the host population density over the last 100 time steps within a patch averaged over all patches, median over 100 replicates. This represents long term host availability. Model parameters of the focal scenario: $\lambda_0 = 4$, intraspecific competition coefficient of the host $\alpha = 0.01$, maximum parasite transmission $\beta_{max} = 8$, shape of trade-off curve $s = 0.5$, searching efficiency of parasite $a = 0.01$.

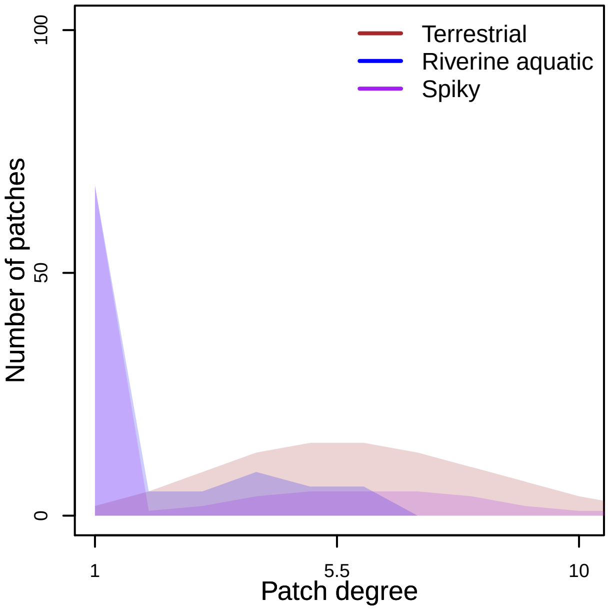


Figure S10: Degree distribution of terrestrial, riverine aquatic landscapes and spiky networks. The x-axis represents patch degree and the shaded region shows the median frequency of a patch of a given degree over 1000 landscape realisations for all landscape types.

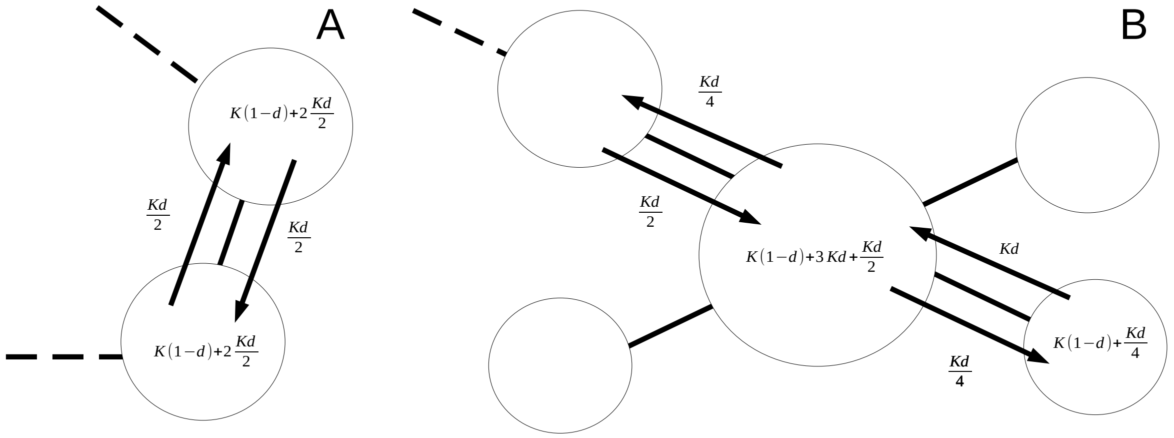


Figure S11: Following Altermatt and Fronhofer (2018), we illustrate how the heterogeneity in distribution of infected densities observed in Fig. 2 are generated. We consider a simplified system in which all patches initially have $K$ individuals, and we assume that in their target patches, each parent produces exactly one offspring. On the patches, we show the number of individuals that are present after one time step. We consider A: a homogeneous landscape in which each patch is connected to a fixed number of patches and B: a heterogeneous landscape. In the homogeneous landscape, the number of dispersers leaving a patch is equal to those entering. However, in heterogeneous networks, patches with a larger degree connected to patches with a lower degree receive more dispersers than they send out, and patches with low degree connected to patches with a higher degree send out more dispersers than they receive. While we just show densities after one time step, these differences between low and high connectivity patches are expected to amplify with time.

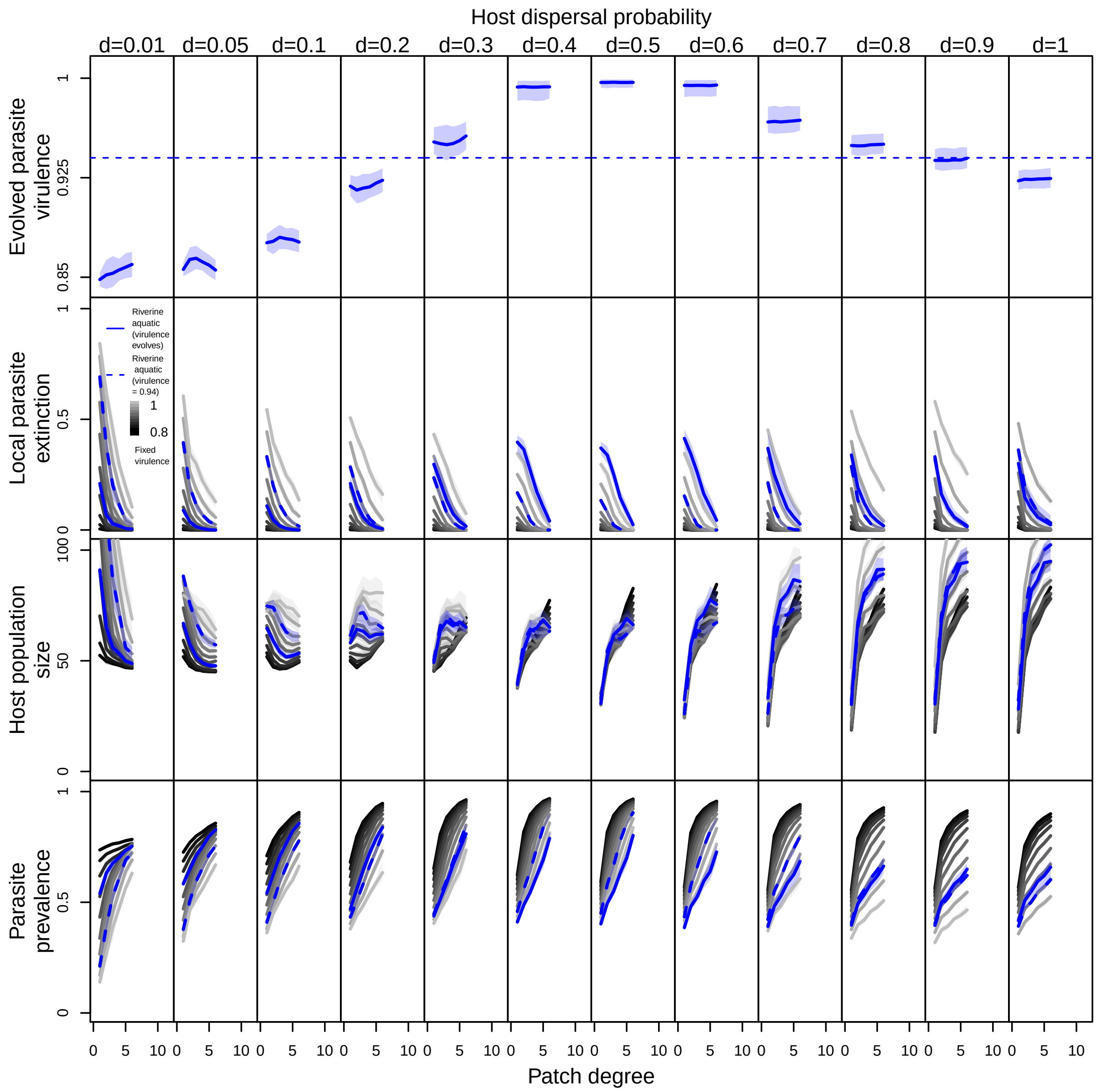


Figure S12: From top to down: spatial distribution of parasite virulence, local parasite extinction, host population size and parasite prevalence as a function of patch degree (number of neighbours of a patch) in riverine aquatic landscapes for fixed ($v = 0.8, ..., 1$; black to grey; fixed virulence value $v = 0.94$ is indicated by blue dashed lines) and evolving virulence (blue solid line). From left to right dispersal increases ($d = 0.01, 0.05, 0.1, 0.2, 0.3, 0.4, 0.5, 0.6, 0.7, 0.8, 0.9, 1$). For all plots patches of the same degree across all landscape realisations are pooled, medians and quartiles are taken over for 100 and 1000 realisations for ecological and evolutionary simulations respectively. Model parameters of the focal scenario: intrinsic growth rate of the host $\lambda_0 = 4$, intraspecific competition coefficient of the host $\alpha = 0.01$, maximum parasite transmission $\beta_{max} = 8$, shape of trade-off curve $s = 0.5$, searching efficiency of parasite $a = 0.01$.

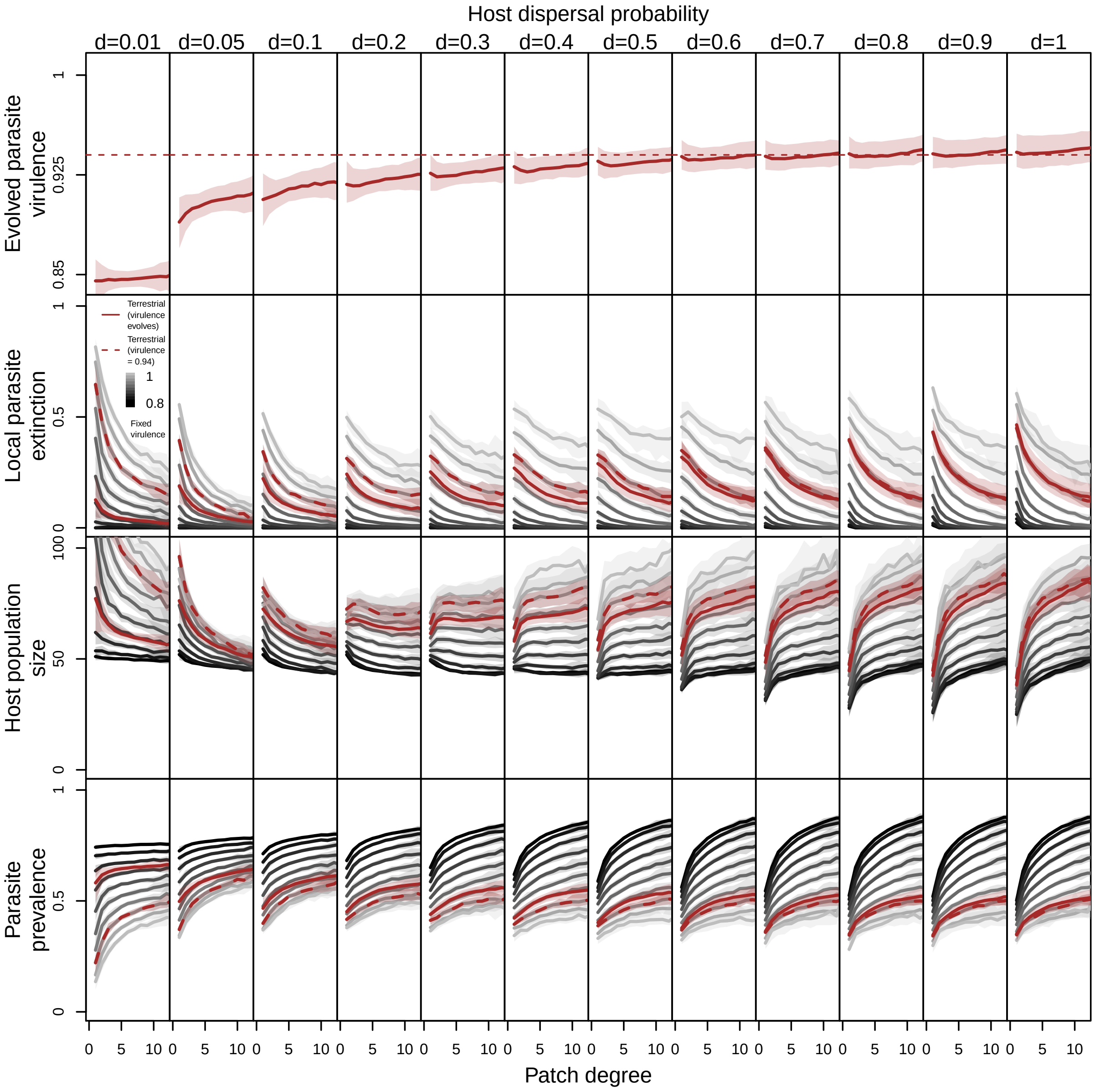


Figure S13: From top to down: spatial distribution of parasite virulence, local parasite extinction, host population size and parasite prevalence as a function of patch degree (number of neighbours of a patch) in terrestrial landscapes for fixed ($v = 0.8, 0.82..., 1$; black to grey; fixed virulence value $v = 0.94$ is indicated by brown dashed lines) and evolving virulence (brown solid line). From left to right dispersal increases ($d = 0.01, 0.05, 0.1, 0.2, 0.3, 0.4, 0.5, 0.6, 0.7, 0.8, 0.9, 1$). For all plots patches of the same degree across all landscape realisations are pooled, medians and quartiles are taken over for 100 and 1000 realisations for ecological and evolutionary simulations respectively. Model parameters of the focal scenario: intrinsic growth rate of the host $\lambda_0 = 4$, intraspecific competition coefficient of the host $\alpha = 0.01$, maximum parasite transmission $\beta_{max} = 8$, shape of trade-off curve $s = 0.5$, searching efficiency of parasite $a = 0.01$.

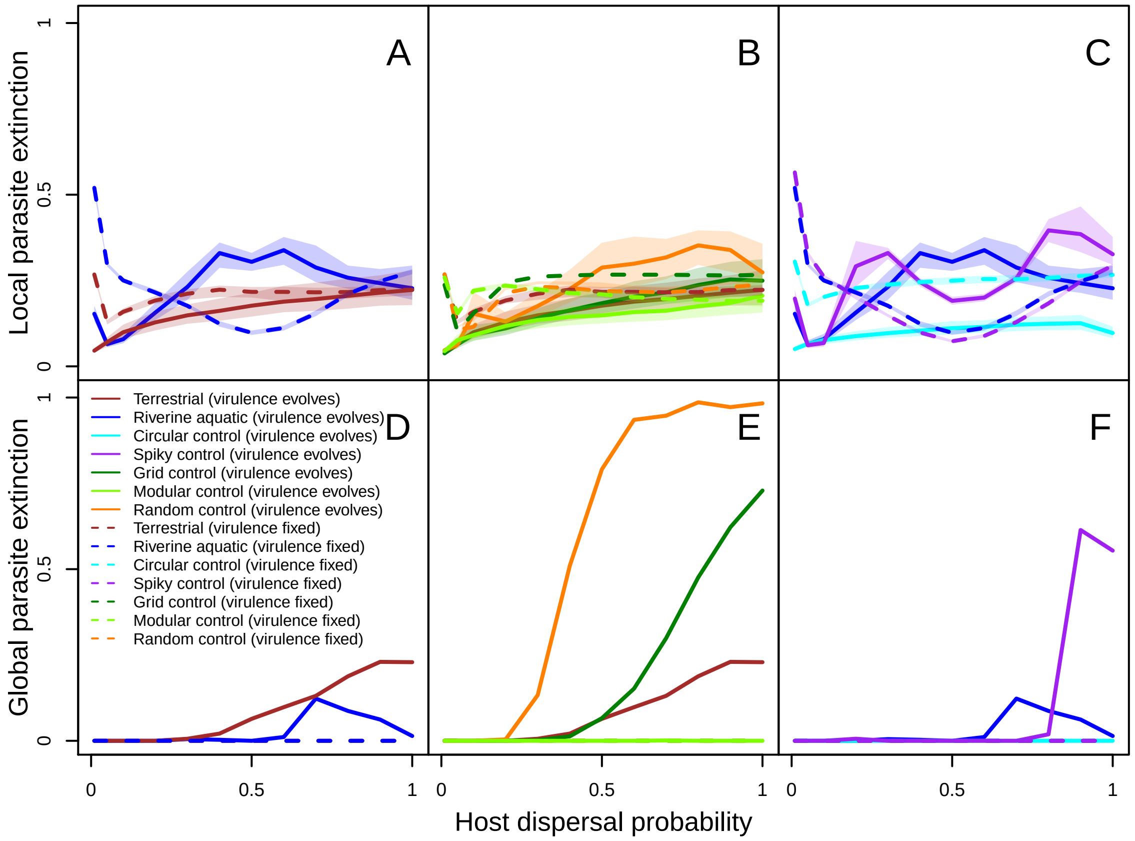


Figure S14: Probability of local (A–C) and global (D–F) parasite extinction as a function of host dispersal ($d = 0.01, 0.05, 0.1, 0.2, 0.3, 0.4, 0.5, 0.6, 0.7, 0.8, 0.9, 1$) in terrestrial vs. riverine aquatic landscapes (A, D), riverine aquatic landscapes vs. controls (B, E) and terrestrial landscapes vs. controls (C, F). A–C: Local parasite extinction as function of host dispersal is calculated as the fraction of patches in which parasite is extinct in the landscape averaged over the last 100 time steps. D–F: Global parasite extinction as a function of host dispersal is calculated as the fraction of realisations in which a parasite goes extinct. The solid lines represent medians of quantities over 1000 realisations of each landscape type in simulations in which virulence evolves and the dashed lines represent the corresponding ecological control for a fixed virulence ($v = 0.94$). In the comparison between terrestrial and riverine aquatic landscapes, we find that while ecologically, local parasite extinctions are predicted to be higher in riverine aquatic landscapes relative to terrestrial landscapes at high and low dispersal rates, and lower at intermediate dispersal rates, virulence evolution leads to greater parasite extinction in riverine aquatic landscapes across dispersal rates. Interestingly, since these parasite extinctions are mostly located in degree-1 patches in the riverine aquatic landscapes, greater parasite extinction locally still leads to fewer scenarios in which the parasite goes extinct globally. The comparison between the terrestrial landscapes and controls shows that while all the landscapes types lead to similar rates of local parasite extinction, random networks lead to the highest rate of parasite extinction globally, followed by grids, terrestrial landscapes and then modular networks. This indicates that both spatial structure and modularity prevent global parasite extinction. Finally, local parasite extinction is lower in circular networks than the heterogeneous networks (riverine aquatic and “spiky networks”) due to evolution of lower virulence which also leads to lower rate of global parasite extinctions in circular networs. Model parameters of the focal scenario: intrinsic growth rate of the host $\lambda_0 = 4$, intraspecific competition coefficient of the host $\alpha = 0.01$, maximum parasite transmission $\beta_{max} = 8$, shape of trade-off curve $s = 0.5$, searching efficiency of parasite $a = 0.01$.

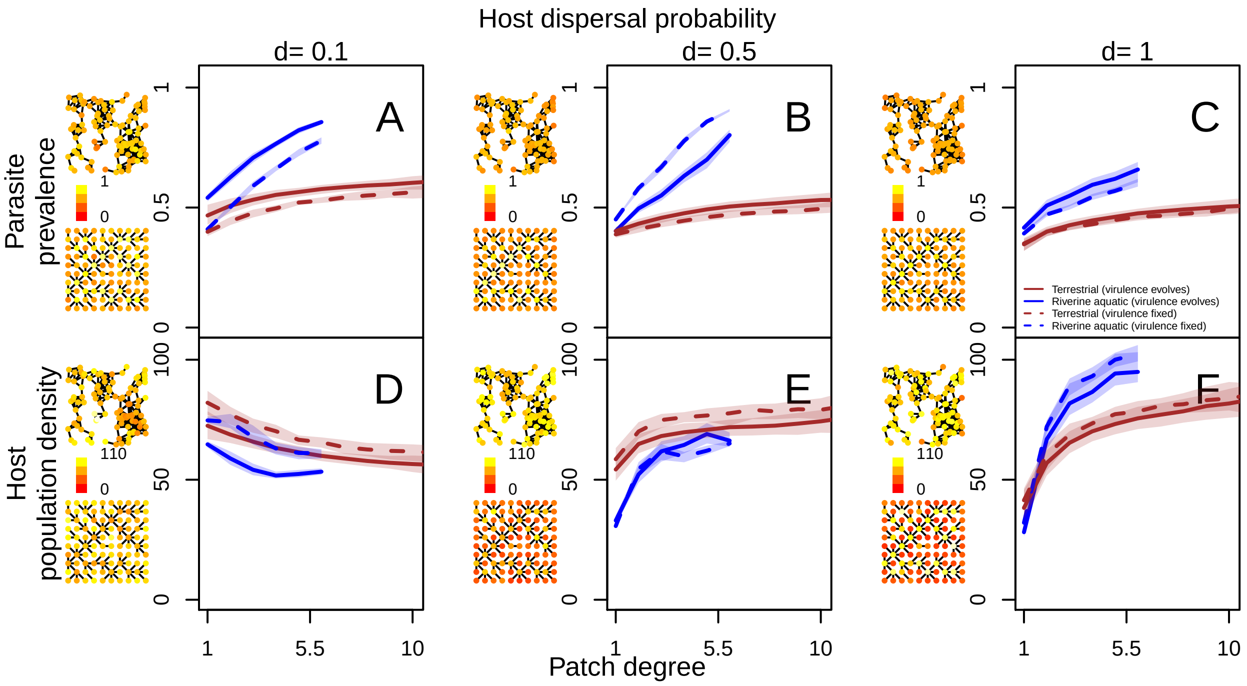


Figure S15: Spatial distribution of parasite prevalence (A–C) and host population density (D–F) as a function of patch degree for terrestrial (brown) and aquatic (blue) landscapes for different host dispersal probabilities ($d = 0.1, 0.5, 1$) in ecological (virulence $v = 0.94$, dashed line) and evolutionary (solid lines) simulations. From left to right, dispersal probability increases. A–C: Parasite prevalence (number of infected individuals divided by the total number of individuals) in a patch, averaged over the last 100 time steps is is plotted as a function of patch degree. On the left side of each plot, the distribution of parasite prevalence in a patch averaged over the last 100 time steps for one representative terrestrial and aquatic landscape when virulence evolves is shown with more yellow colours indicating higher prevalence. D–F: Host population density averaged over the last 100 time steps is plotted as a function of patch degree. On the left side of each plot, the distribution of average host population density when virulence evolution is taken into account over the last 100 time steps for one representative terrestrial and aquatic landscape is shown with more yellow colours indicating higher average host population density. For all plots patches of the same degree across all landscape realisations are pooled, medians and quartiles are taken over for 100 and 1000 realisations for ecological and evolutionary simulations respectively. Predictions based on fixed virulence of spatial distribution of parasite prevalence and host population density remain relatively unchanged compared to when evolution of virulence is taken into account. Model parameters of the focal scenario: intrinsic growth rate of the host $\lambda_0 = 4$, intraspecific competition coefficient of the host $\alpha = 0.01$, maximum parasite transmission $\beta_{max} = 8$, shape of trade-off curve $s = 0.5$, searching efficiency of parasite $a = 0.01$.
